# Cellulose fermentation by the gut microbiota is likely not essential for the nutrition of millipedes

**DOI:** 10.1101/2024.03.01.582937

**Authors:** Julius Eyiuche Nweze, Shruti Gupta, Michaela M. Salcher, Vladimír Šustr, Terézia Horváthová, Roey Angel

## Abstract

Millipedes are thought to depend on their gut microbiome for processing plant-litter-cellulose through fermentation, similar to many other arthropods. However, this hypothesis lacks sufficient evidence. To investigate this, we disrupted the gut microbiota of juvenile *Epibolus pulchripes* (tropical, CH_4_-emitting) and *Glomeris connexa* (temperate, non-CH_4_-emitting) using chemical inhibitors and isotopic labelling. Feeding the millipedes sterile or antibiotics-treated litter notably reduced faecal production and microbial load without major impacts on survival or weight. Bacterial diversity remained similar, with *Bacteroidota* dominant in *E. pulchripes* and *Pseudomonadota* in *G. connexa*. Sodium-2-bromoethanesulfonate treatment halted CH_4_ emissions and reduced the faecal *mcrA* copies in *E. pulchripes* after 14 days, but emissions resumed after returning to normal feeding. Methanogens in the order *Methanobacteriales* and *Methanomasscilliicoccales* associated with protists were detected using Catalysed Reporter Deposition Fluorescence *In situ* Hybridization (CARD-FISH) on day 21, despite suppressed CH_4_-emission. Employing ^13^C-labeled leaf litter and RNA-SIP revealed a slow and gradual prokaryote labelling, indicating a significant density shift only by day 21. In addition to labelling of taxa from orders well-recognized for their role in (ligno)cellulose fermentation (e.g., *Bacteroidales*, *Burkholderiales*, and *Enterobacterales*), others, such as members of *Desulfovibrionales* were also labelled. Surprisingly, labelling of the fungal biomass was somewhat quicker. Our findings suggest that fermentation by the gut microbiota is likely not essential for the millipede’s nutrition.

**Importance:** Millipedes (Diplopoda) constitute the third most significant group of detritivores after termites and earthworms, yet they have been comparatively understudied. Traditionally, it was believed that millipedes gain energy from fermenting cellulose using their gut microbiota, similar to wood-feeding termites, but this belief lacks evidence. This study used two model millipede species, *Epibolus pulchripes* (large, tropical, and methane emitter) and *Glomeris connexa* (small, temperate, and non-methane emitter) to test this belief. We used chemical manipulation experiments, stable isotope labelling, and DNA sequencing to comprehend the microbiota’s role in the millipede’s nutrition. The findings suggest that cellulose fermentation by the gut microbiota may not be essential for millipede nutrition; instead, bacteriovory and fungivory might be the dominant feeding strategies of millipedes.

## Introduction

Like most animals, invertebrates form intricate partnerships with diverse microbial communities (1), contributing significantly to their evolutionary and ecological success (2). This close interconnectedness has led to the concept of animals as “holobionts,” where the host and its microbiota are viewed as a single ecological entity (3, 4). Recent studies on microbiomes provide further evidence of the widespread prevalence of microbial partnerships across the animal kingdom (5, 6).

While most invertebrates have microbial associations, their reliance on them varies widely. Termites, for instance, depend entirely on their gut microbiota for nutrition (7). Conversely, many other arthropods, such as caterpillars, may lack a resident gut microbiota and develop fully even germ-free (8). Most arthropods generally fall between these extremes, relying on their microbiota for some form of support (e.g. cockroaches (9, 10) or isopods (11, 12)). Detritivorous and xylophagous animals typically rely on gut microorganisms for cellulose digestion. Although animal cellulases are found in some gut systems (13), (ligno)cellulolytic bacteria, fungi and protists are generally deemed necessary for hydrolysis and fermentation, releasing short-chain fatty acids, which get absorbed by the host (14).

Millipedes (Diplopoda) are crucial detritivores widely distributed and abundant in many temperate and tropical ecosystems (15). Despite their status as keystone species in tropical and temperate forests (16), millipedes have been understudied compared to other detritivores, particularly concerning their microbiome. Due to the nutrient-poor nature of plant litter, millipedes compensate for low assimilation efficiencies through high ingestion rates (17). Similar to other arthropods, millipedes host diverse gut microorganisms (18). Notably, the central hindgut was shown to host the highest microorganism density, attaching to its cuticle, while the foregut and midgut contain mostly transient inhabitants (19). Various studies suggest that certain millipede gut bacteria possess enzymes for breaking down plant polysaccharides (20–24). If millipedes rely on cellulose for their nutrition, extensive fermentation followed by methanogenesis, similar to ruminants or wood-feeding termites, should occur in their guts (7, 25). However, methanogenesis has only been observed in some millipede species, but not others, with its occurrence correlated to the millipede size (26). Despite these findings, direct proof of gut microorganisms supporting the millipede’s nutritional needs has not been experimentally demonstrated. An alternative hypothesis suggests millipedes foster microbial growth in litter, potentially digesting the resulting fungal and bacterial biomass (27).

To investigate the role of the millipede gut microbiota, we conducted experiments using two model species: the CH_4_-emitting *Epibolus pulchripes* (Spirobolida) and *Glomeris connexa* (Glomerida), which do not emit CH_4_. *E. pulchripes* is a large millipede (130–160 mm) common along the East African coast (28), while *G. connexa* is smaller (10-17 mm) and native to Central Europe (29). We assessed the impact of inhibitors on body weight, survival, faecal bacterial load, gut bacterial composition, and CH_4_ production. Additionally, we identified metabolically active hindgut prokaryotes using ^13^C-RNA-SIP.

## Results

### Effect of antibiotic curing

Feeding millipedes with either sterile or antibiotics-treated feed led to only negligible weight change in both species (Fig 1a and b; Table S2) with no significant trend. The treatment also did not significantly impact the millipedes’ survival based on Kaplan-Meier estimates (Fig. S1). Despite maintaining a stable weight, faecal production decreased over time in response to antibiotics or sterile feed (*P < 2.2e-16* for both species; Fig. 1c and d; Table S3). No significant difference was found between the treated groups in *E. pulchripes*, but in *G. connexa* the sterile-litter group was different from the antibiotic-treated groups (*P < 0.0001*). Total faecal colony counts in both millipede species were also consistently higher in the control group compared to the antibiotic-treated or sterile feeding groups at all time points (*P < 0.0001*; Fig. 1e and f; Table S4). After 35 days for *E. pulchripes* and 16 days for *G. connexa*, most animals in the treatment groups ceased faecal production, leading to the cessation of plate counts. Once again, only the sterile-litter group in *G. connexa* differed from the other treatment groups. Total faecal 16S rRNA gene copies in *E. pulchripes* were reduced by 61%–77% in the treated groups compared to the control group (*P = 0.01*), while In *G. connexa*, 34%–74% reductions were observed in the treated groups (*P < 0.001*; Fig. 1g; Table S5). In both species, no difference between the treated groups was observed. After noting a substantial decrease in bacterial load, we measured CH_4_ emission on day 35 (Fig. 1h; Table S6). As anticipated, CH_4_ was present in *E. pulchripes* but absent in *G. connexa* (data not shown). The control groups displayed a significantly higher CH_4_ production rate (284.1 ± 58 nmol mg^-1^ d^-1^) than the other treatments (P = 0.0008). However, the treated groups saw a 57‒74% reduction in CH_4_ production without significant differences between them.

**Fig. 1.**
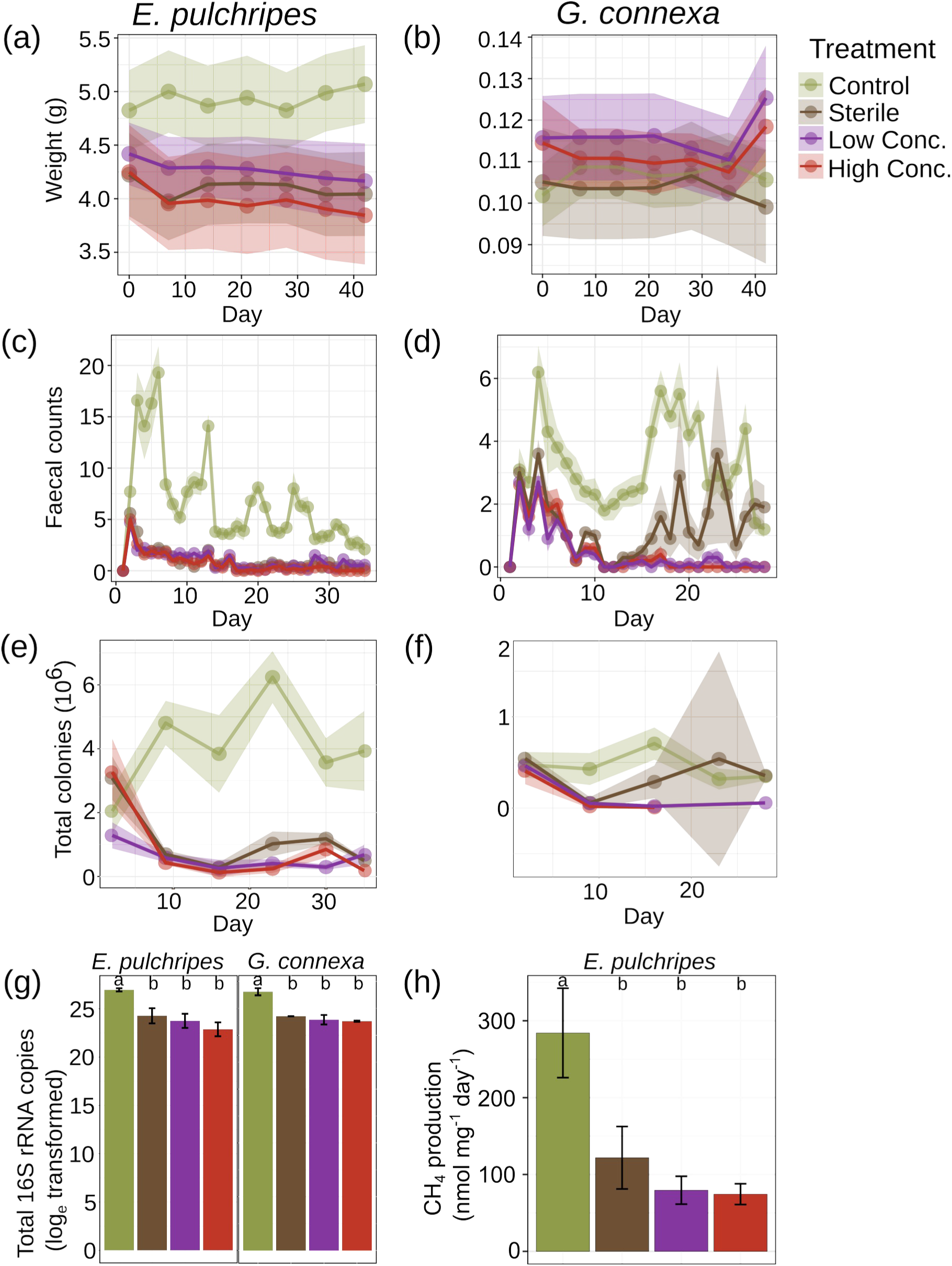
Effect of antibiotic treatment on *E. pulchripes* and *G. connexa*. Time series of mean weight loss (mean ± SE ribbon) in (a) *E. pulchripes* and (b) *G. connexa*; faecal counts in (c) *E. pulchripes* and (d) *G. connexa*; total colony forming units in (e) *E. pulchripes* and (f) *G. connexa*; (g) 16S rRNA gene copy numbers in the faeces; and (h) CH_4_ production rate after 35 days of antibiotics treatment in *E. pulchripes*. ’High Conc.’ and ’Low Conc.’ refer to the concentration of applied antibiotics (see Materials and Methods for more details). Different lower case letters in panels g and h denote statistical significance. See Results for a detailed description of the statistical tests performed on the time series (panels a-f).

### Prokaryotic community compositions after treatment

We sequenced 48 samples of *E. pulchripes* and *G. connexa*, consisting of 12 hindguts and 12 faecal samples for each species. The average sequencing depth stood at ca. 40K reads per sample, post-processing of reads and decontamination (Table S7 and S8). The two millipede species differed remarkably in their microbial composition, with the phylum *Bacteroidota* dominating the hindgut of *E. pulchripes* and *Pseudomonadota* that of *G. connexa*. In each case, these phyla comprised over 50% of the abundance regardless of treatment (Fig. 2a and b; Table S9). *Pseudomonadota*, *Bacteroidota* and *Actinobacteriota* dominated both species’ faecal pellets. (Fig. 2c and d). On the genus level, *E. pulchripes*’ hindgut and faecal samples were primarily dominated by *Citrobacter*, *Bacteroides*, and *Pseudomonas* (Fig. 2e-h; Table S9). In contrast, *G. connexa* showed differences between hindgut and faecal sample compositions, with faecal samples appearing more diverse (Fig. 2h).

**Fig. 2.**
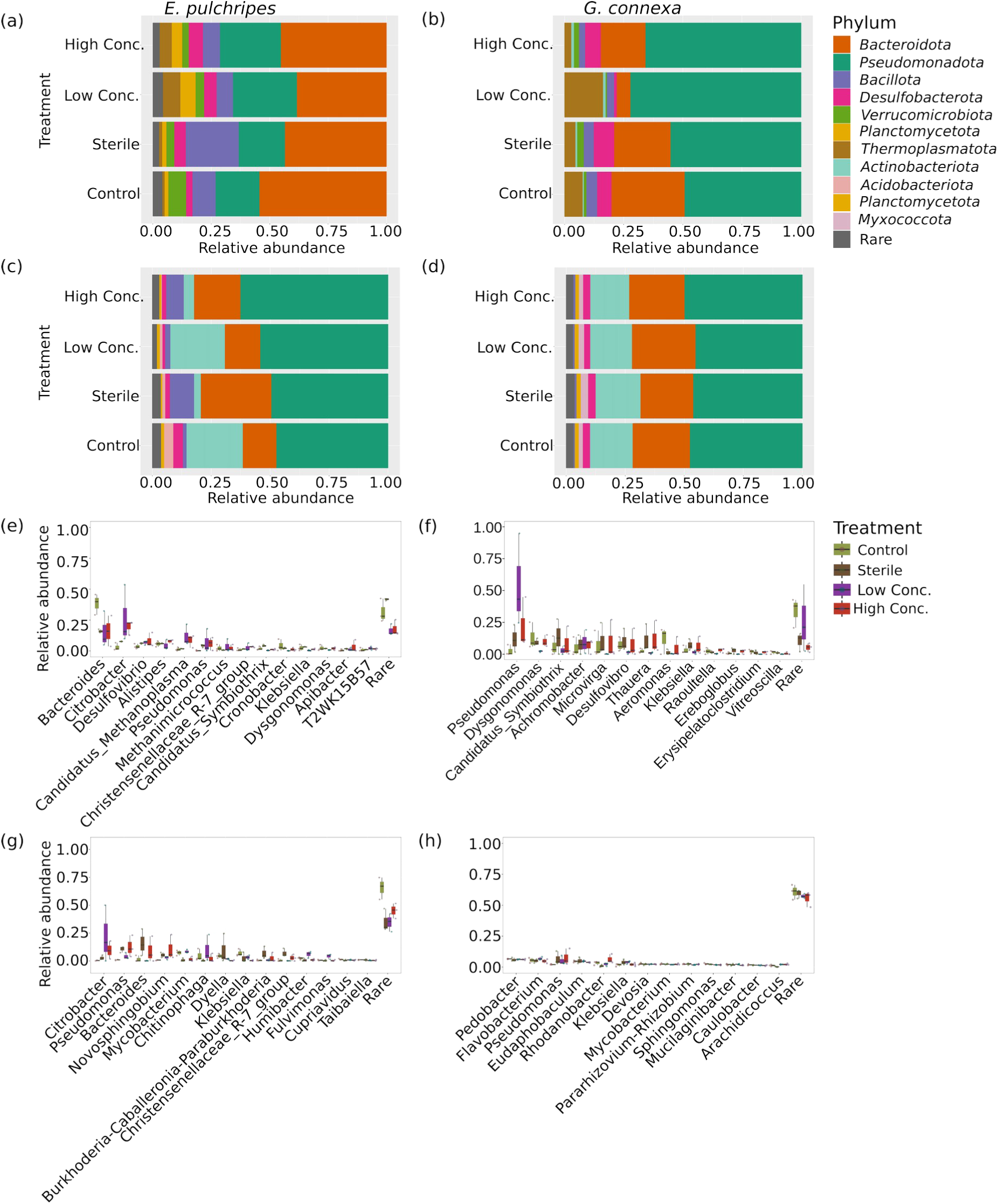
Effect of antibiotic treatment on the taxonomic composition of prokaryotes in *E. Pulchripes* (left) and *G. Connexa* (right) following treatment. Phylum distribution in the hindguts (a and b) and the faeces (c and d). Distribution at genus level in the hindguts (e and f) and faeces (g and h). ’High Conc.’ and ’Low Conc.’ refer to the conc. of antibiotics applied (see Materials and Methods for more details).

### Impact of treatment on prokaryotic community structures

Overall, no significant differences were found in alpha diversity within or between treatment groups in the hindguts (Fig. 3a & b; Fig. S2; Table S10) or faeces (Fig. 3c & d; Fig. S2; Table S10) of *E. pulchripes* and *G. connexa*. *E. pulchripes*’ hindgut groups displayed greater bacterial diversity and richness than *G. connexa*. Constrained analysis of principal coordinates (CAP) revealed significant differences in microbial community composition among sterile feeding or antibiotics treatments in both hindguts and faeces of both species (Fig. 3e, f, g & h; Table S10). ANCOM-BC2 analysis identified only a handful of microbial genera with differential abundance between treatments (Table S11; Fig. S3), indicating that the antibiotic treatment worked relatively non-selective. The few taxa with a decrease in the mean absolute abundance (e.g. *Streptomycetaceae* and *Mucilaginibacter* from the *E. pulchripes*’ faeces) are known to often possess antibiotic resistance genes.

**Fig. 3.**
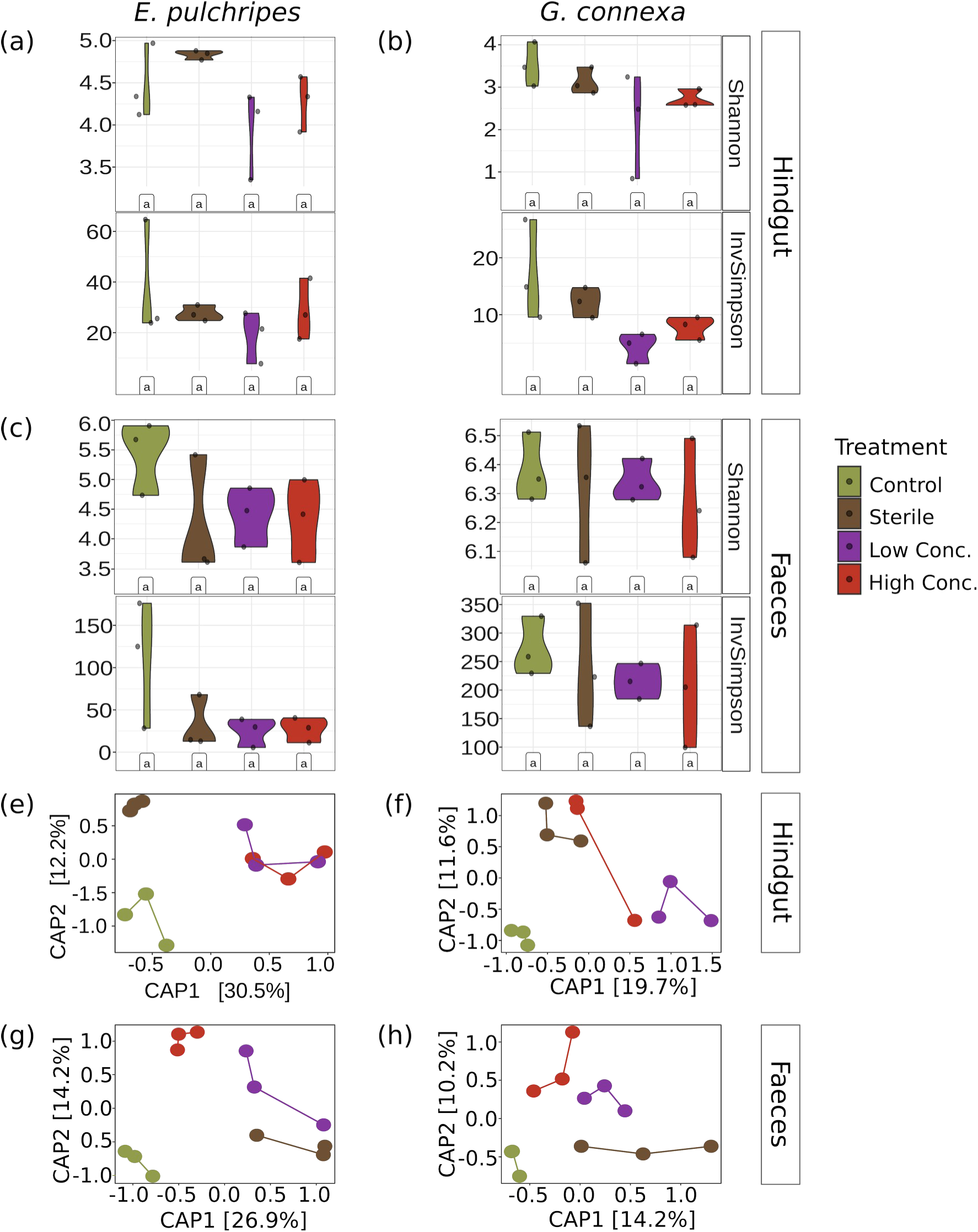
Effect of antibiotic treatment on the alpha and beta diversity indices of the microbial communities in the hindgut and faeces in *E. pulchripes* (left) and *G. connexa* (right). Alpha diversity values for each species, stratified by treatment groups for hindgut (a and b) and faeces samples (c and d) from *E. pulchripes* and *G. connexa*. The statistical test was based on Kruskal–Wallis (identical letters denote p >0.05). Dissimilarity between hindgut (e and f) and faeces (g and h) microbial communities in the different treatments using constrained principal coordinates analysis (PcoA) with the model Dist.Mat ∼ Treatment for each species and sample type separately.

### Influence of BES inhibition on methanogenesis in *E. pulchripes*

Na-BES-treated litter was provided to investigate the importance of methanogenesis in the CH_4_-emitting *E. pulchripes*. Methane emissions showed no significant differences on days 0 (P = 0.19) and 7 (P = 0.08; Fig. 4A; Table S12). However, by day 14, CH_4_ production was nearly fully inhibited (P = 2.7 x 10^-4^) and remained so for an additional 21 days. Upon switching to untreated litter on day 35, CH_4_ emissions began recovering on day 49 and resumed pre-treatment values by day 63. Despite some average weight increase in the treated groups, no significant difference was detected at any time (Fig. 4b).

**Fig. 4.**
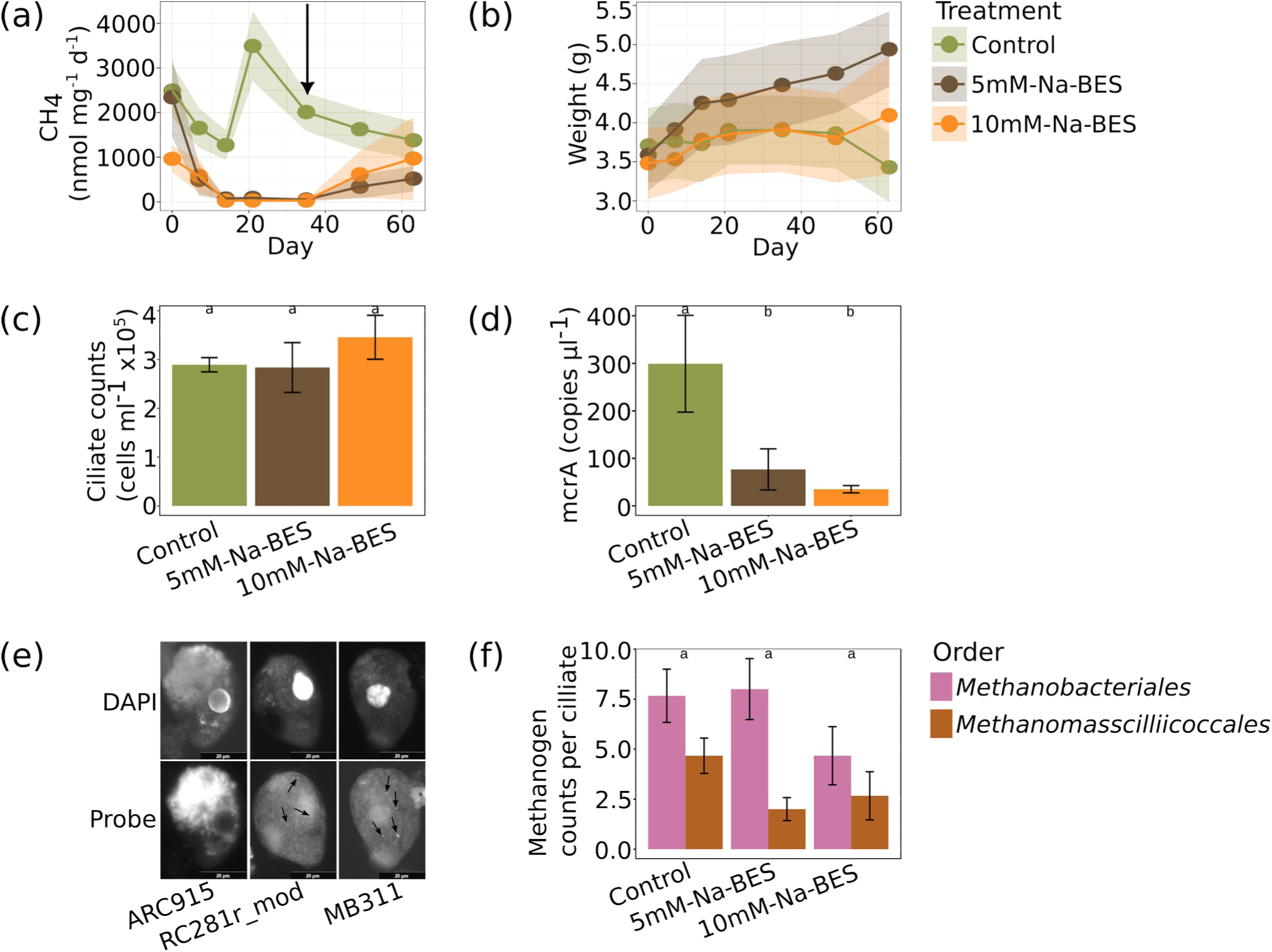
Effect of BES treatment on CH_4_ emissions from *E. pulchripes*, animal weight, ciliates and ciliate-associated methanogens. (a) Emission rates of CH_4_ over time followed by recommence of methane production after the switch to untreated litters (indicated by the arrow). (b) Change in the weight of *E. pulchripes* over time. (c) Enumeration of symbiotic ciliates found in the faeces following BES treatment. (d) *mcrA* gene copy numbers in the faecal samples following BES treatment. (e) Fluorescence microscopy images of ciliates and the two most-abundant endosymbiotic methanogens in faecal samples of *E. pulchripes* using DAPI and CARD-FISH probes. ARC915: general archaea, RC281r_mod: *Methanomassciillicoccales*, and MB311: *Methanobacteriales* in the 10mM-Na-BES-treated group. (f) Enumeration of the methanogens associated with ciliates using FISH signals.

After inhibiting methane production for 21 days, a suspension made from fresh faeces was examined under a bright-field microscope, revealing various protists, nematodes, and rotifers ranging from 12 to 100 μm in size (Fig. S4). The ciliate abundance averaged 3 × 10^5^ ml^-1^, regardless of treatment (Fig. 4c; Table S13). Quantification of the *mcrA* gene, pivotal in methanogenesis (30), showed a significant reduction in the two Na-BES-treated groups compared to the control (P = 0.02; Fig. 4d; Table S13). CARD-FISH was used to detect the presence of free-living (Fig. S5) and symbiotic archaea (Fig. S6), primarily methanogens, in protists from faecal samples. The amplicon sequencing data indicated that members of the *Methanomassciillicoccales* and *Methanobacteriales* were the dominant methanogens in *E. pulchripes*, and these orders were accordingly targeted. Although *mcrA* copy numbers declined, positive hybridisation signals for these methanogen orders were observed in both Na-BES treatments. Methanogens were detected on the 0.2 µm filter (Fig. S5) and associated with protists as endosymbionts (Fig. 4e; Fig. S6), with no significant changes in its count per ciliate (Fig. 4f).

### Detection of active microbiota with ^13^C-RNA-SIP

RNA-SIP was used to identify the active microorganisms in the millipedes’ gut on a temporal scale (Table S14). The shift in peak of 16S rRNA towards the denser gradient fractions, indicating label incorporation, was evident by day 3 and more prominently by day 7 for *E. pulchripes* and day 14 for *G. connexa* (Fig. 5). Nevertheless, despite feeding on fully-labelled litter for 21 days, a significant portion of RNA remained unlabelled. Surprisingly, the labelling of the fungal biomass, represented by the 18S rRNA peak, shifted earlier towards denser gradient fractions compared to 16S rRNA in both millipede species (Fig. S7). However, the lack of pronounced peak deviation compared to the control in some replicates and days does not necessarily imply unsuccessful labelling since the labelled fraction of the community might still be too small. Indeed, there was a significant change in community composition in the heavy fractions of labelled gradients compared to unlabelled ones already by day 3 (Fig. S8; Table S15).

**Fig. 5.**
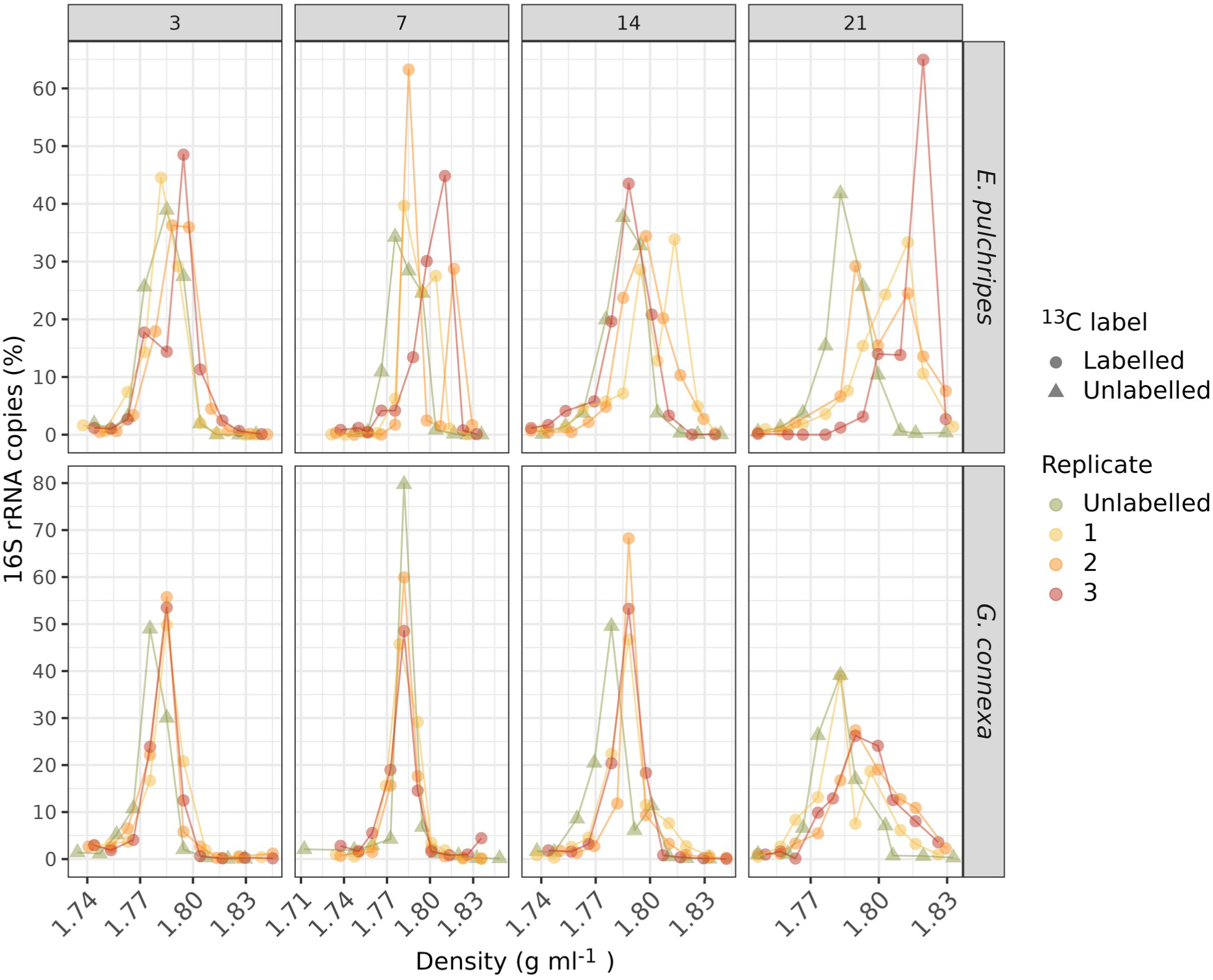
Bacterial 16S rRNA copies recovered from each fraction in the SIP gradients. rRNA copies relative to the total number of rRNA copies obtained from the entire gradient against the buoyant density of each fraction. Labelled RNA is expected to be found in fractions with density >1.795 g ml^-1^.

For comparing heavy fractions in labelled versus unlabelled gradients of 16S RNA, an average of 1305 ± 59 and 579 ± 41 ASVs were used for *E. pulchripes* and *G. connexa* per time point after filtering (Table S16). Surprisingly, the model identified, on average, only 22% of the ASVs in *E. pulchripes* and 24% in *G. connexa* as labelled. These values were consistent over time. Therefore, the shift in copy-number peaks towards denser fractions, as observed in Fig. 5, was due to increased labelling in already labelled ASVs rather than a change in the proportion of labelled ASVs.

### Diversity of active microbiota in a heavy fraction of ^13^C-RNA-SIP

In agreement with the general bacterial diversity in the gut, the major phyla whose members were flagged as labelled were *Actinobacteriota, Bacillota*, *Bacteroidota*, and *Pseudomonadota* (Fig. 6; Table S16). In *E. pulchripes*, *Bacillota* comprised 35 to 55.3%, *Bacteroidota* 13.1 to 15.1% and *Pseudomonadota* from 13.8 to 23% of the total labelled ASVs. In *G. connexa*, *Bacillota* comprised 20.4 to 45.9% of total significant ASVs, *Pseudomonadota* ranged from 20 to 51.6%, *Actinobacteriota* from 15.1% to 22.6%, and *Bacteroidota* from 3.2 to 10.8%. Fig. S9-15 show the phylogenetic distribution of the labelled ASVs across the samples in each of the major bacterial classes. Despite our expectation for gradual labelling of the microorganisms with time, similar numbers and, in many cases, the same ASVs were consistently labelled throughout the incubation. In *E. pulchripes,* members of the classes *Clostridia* and the orders *Bacteroidales, Rhizobiales*, *Enterobacterales*, *Desulfovibrionales*, *Pirellulales*, *Verrucomicrobiales* and *Victivallales* were most prominently labelled. In *G. connexa*, members of the class *Clostridia* and the orders *Bacteroidales, Rhodobacterales*, *Enterobacterales*, *Pseudomondales* and Micrococcales were most prominently labelled.

**Fig. 6.**
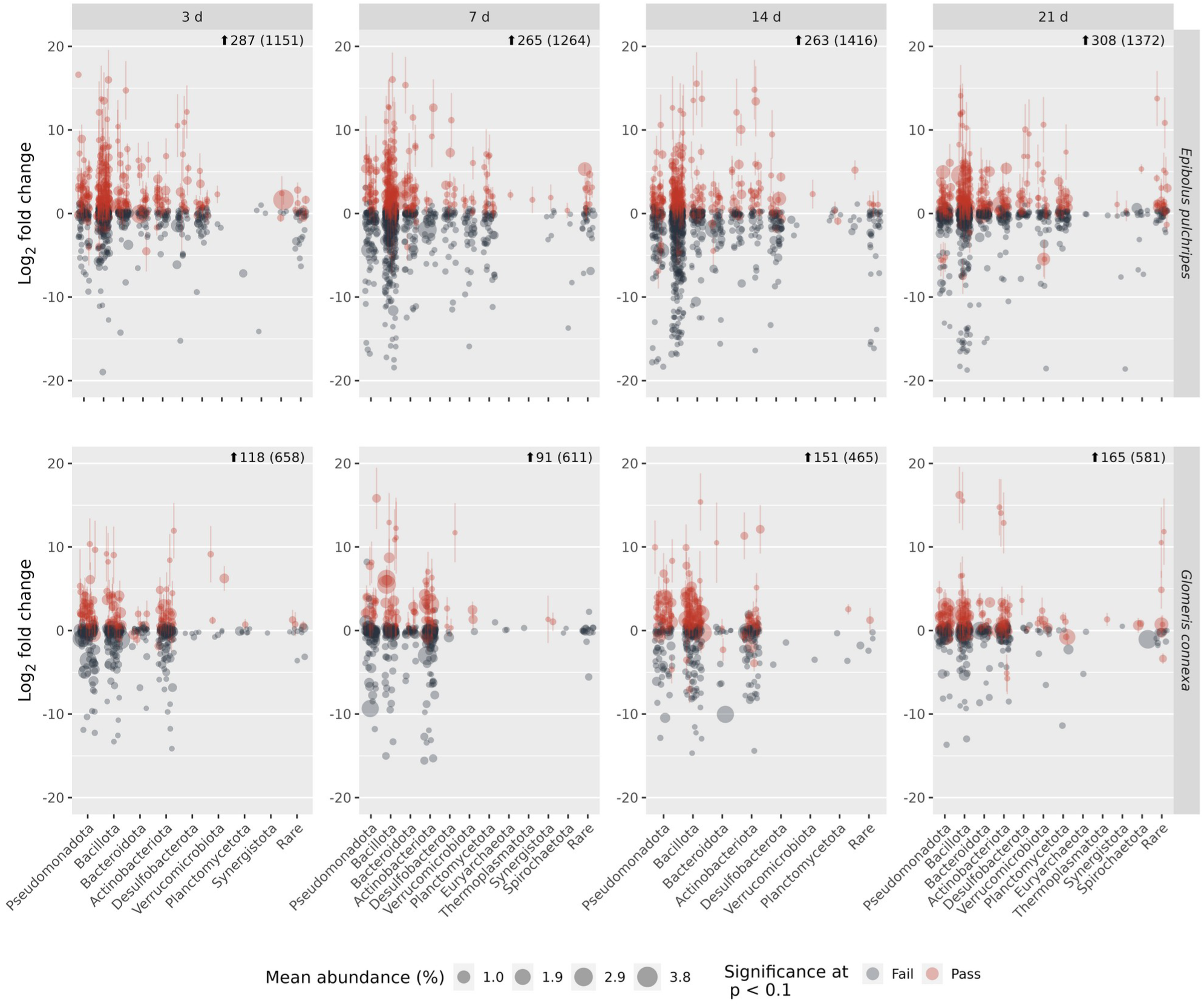
Differentially abundant ASVs between the labelled and unlabelled gradients of the SIP experiments. Comparison of the relative abundance of each ASV from *E. pulchripes* and *G. connexa*. Each subfigure represents a triplicate. The plot shows the most abundant phyla in the dataset in decreasing abundance. The differential abundance of any particular ASV is given in Log_2_ fold change. “Rare” indicates phyla with mean relative abundance below 0.45%.

## Discussion

The gut microbiota, crucial for the ecophysiology of arthropods (31), is especially vital for detritivores relying on recalcitrant plant polymers with low nitrogen content. Building on culture-based (32) and recent molecular studies (21, 23, 33, 34), the findings underscore a generally stable and species-specific millipede gut microbiota, resistant to inhibitors. Variations in closely related arthropods may arise from gut conditions like pH, oxygen availability (35), and gut topography (19). Specifically for millipedes, hindgut volume differences, influencing redox potential, likely contribute to microbiota variations, promoting fermentation and methanogenesis in larger species (e.g., *E. pulchripes* and *T. aoutii*) but not in smaller ones (e.g., *G. connexa*) (21, 26).

Curing or sterilizing arthropods to assess their dependence on gut microbiota has been conducted in various species, yielding diverse outcomes. Not surprisingly, for wood-feeding termites, exposure to high oxygen levels results in the disappearance of flagellates, leading to starvation (7, 36). This is because wood-feeding termites rely on short-chain fatty acids, which are the products of cellulose fermentation, for their nutrition. Cured arthropods in other studies exhibited moderate responses, including decreased feeding and altered microbiota, observed in desert millipedes (24), Carabidae members (37), and egg-hatching cockroaches (38). In contrast, larval Lepidoptera, exclusively feeding on fresh leaves and likely relying on simple sugars, showed no physiological response to antibiotic curing (39). Both millipede species in this study maintained a stable weight, suggesting they might not require fermentation products for nutrition. However, the notable decrease in faecal production and the relatively unchanged taxonomic composition indicated a potentially significant role in the microbiota. Notably, there was a shift in abundance towards antibiotic-resistant bacterial strains, such as *Citrobacter* and *Bacteroides* in *E. pulchripes* (40, 41) and *Pseudomonas* and *Achromobacter* in *G. connexa* (42, 43).

This study validated CH_4_ release in *E. pulchripes*, aligning with previous findings (26, 44). Antibiotics decreased CH_4_ emission, likely disrupting bacterial fermentation, a phenomenon observed in cockroaches when bacteria and flagellates were targeted (45). As expected, the application of BES, a specific methanogenesis inhibitor (46), reduced CH_4_ production to undetectable levels without apparent effects on *E. pulchripes* fitness. As CH_4_ production serves as a hydrogen sink in anaerobic systems driving syntrophic fermentation processes (47), it supports the notion that gut fermentation is non-essential for millipede nutrition. The dominant methanogens, *Methanobacteriales* and *Methanomassiliicoccales*, in our millipedes are known gut inhabitants (44). Surprisingly, despite suppressed CH4 production and a 10-fold drop in mcrA gene copy numbers, methanogen density in the gut remained unaffected. In dynamic gut systems, members must continue to proliferate to avoid being flushed out, methanogens likely live as symbionts of protists, directly benefiting from fermentation products, similar to the case in termites (48, 49).

In the SIP experiment, RNA labelling was slow and gradual, leaving a substantial portion unlabelled even after 21 days, indicating the inefficiency of the millipede gut system in degrading leaf litter and assimilating carbon. In contrast, fungal biomass exhibited faster and higher labelling, especially in G. connexa. Soil litter decomposition studies suggest fungi thrive first on recalcitrant and nutrient-poor litter, with bacteria flourishing later on nutrient-rich and more labile litter (50, 51). In the hindgut of both millipede species (21) and *Telodeinopus aoutii* (23), *Ascomycota* and *Basidiomycota* dominate, mirroring soil decomposition patterns (50, 52, 53).

Despite millipedes’ ability to hydrolyze polysaccharides, lipids, and proteins through salivary gland enzymes alongside their resident microbes (as in many other detritivores; 13, 54, 55) and conditions, and methanogenesis in the digestive tract (26, 44, 56), cellulose digestion significance in millipede metabolism remains inconclusive. Quantitative data, including low metabolic rates in millipedes fed pure cellulose, suggest challenges in maintaining a positive energy balance (57).

The labelled microbiota in *E. pulchripes* and *G. connexa*, primarily *Bacillota*, *Bacteroidota*, and *Pseudomonadota*, show distinctive patterns associated with polysaccharide degradation, consistent with recent millipede studies (21, 23). Similar labelling of these phyla was observed in a scarab beetle study using 13C-cellulose (58). Although certain labelled taxa (e.g., *Bacteroidales*, *Burkholderiales*, and *Enterobacterales*) are recognized for their role in (ligno)cellulose fermentation in millipedes (21, 23, 34), others (e.g., members of *Desulfovibrionales*) are hindgut microorganisms involved in processes like sulfate reduction and are likely unrelated to fermentation. Despite senescent leaves not being exclusively comprised of (ligno)cellulose, these polymers constitute approximately 50–75% of litter material (59). In the near absence of other terminal electron acceptors in the gut, most other simpler carbon sources will also need fermentation for metabolism. Consequently, we conclude that while cellulolytic fermentation occurs in the millipede gut, it likely contributes minimally to the host’s diet.

If fermentation products are not a primary nutritional source for the millipede, their main nutrient origin remains a question. Classical ^14^C-labelling studies indicated bacterial assimilation into the millipede’s biomass surpassing that of plants but focused on lab-grown strains and omitted fungi (27). Woodlice, another detritivore, exhibit a preference for fungi- or bacteria-colonized leaf tissues over natural litter (60, 61). Genomic and transcriptomic screening of the studied millipede species revealed glycoside hydrolases (GH) capable of degrading chitin and peptidoglycan as abundant as, or even more so than, cellulose-degrading GHs (21). The decrease in ergosterol levels post-digestion supports significant fungal digestion in the millipede gut (62). Some species exhibit a preference for fungal fruiting bodies, algae, and lichen films (63). Millipedes’ midgut fluid effectively kills bacteria in a species-specific manner (64). Coprophagy in millipedes may provide access to fresh microbial and fungal biomass resulting from a partial breakdown of recalcitrant plant material (65). Additionally, millipedes produce endogenous GHs in their salivary glands and midgut for digesting non-structural plant material (23, 34, 55). Fluid feeding, described in Colobognatha millipedes, enables feeding on fresh plant material (66). These findings don’t exclude other roles of the millipede gut microbiota, such as detoxification of plant toxins (67), essential compound production (23), protection against pathogens (33), and even acquiring new genes through horizontal transfer (68).

This work demonstrates that cellulose fermentation likely plays a minor role, at best, in the millipede’s nutrition. Further work is needed to decipher their exact trophic function in nature and the potential role their microbiota plays in their survival and modulating greenhouse gas emissions.

## Materials and Methods

### Animal collection and maintenance

We used juvenile *E. pulchripes* from our lab breeding colony and wild-caught *G. connexa* from Czechia (forest near Helfenburk u Bavorova; 49°8’10.32“ N, 14°0’24.21” E). No specific permit was required for the collection. Species identification relied on morphological features (69, 70); data not shown). Before use, the animals were kept in the lab for several weeks. Both species were housed in perforated plastic terraria, filled with commercial sand as a substrate, broken terracotta pots for shelter, and locally collected or purchased Canadian poplar (*Populus x canadensis*) leaf litter (see below). Moisture (50-60%) was maintained by spraying with tap water every other day. Both species experienced a 12-hour photoperiod. *E. pulchripes* were housed individually in a box (19.3 x 13.8 x 5 cm) at 25 **°**C and in a climate-controlled room. Meanwhile, five *G. connexa* individuals were kept in each box (15 x 10 x 4 cm) in an incubator (TERMOBOX LBT 165, Vanellus s.r.o.) at a temperature of 15 **°**C.

### Antibiotic curing

Each millipede species comprised 40 individuals split into four groups of ten: Control, Sterile, diluted antibiotics (2X-Diluted in *E. pulchripes* and 5X-Diluted in *G. connexa*) and undiluted antibiotics (Undiluted in *E. pulchripes* and 2X-Diluted in *G. connexa*). Briefly, the Control group was fed untreated, senesced leaves, the Sterile group was fed autoclaved leaves, and the antibiotics-treated groups were fed autoclaved leaves treated with antibiotics. *E. pulchripes* groups were fed around 2.4 g of litter, while *G. connexa* groups received 0.5 g. Just before feeding, the leaf litter was sprayed with 500 µl of tap water (Control), sterile distilled water (Sterile), or antibiotics solution containing penicillin G: 10,000 units ml^-1^, streptomycin sulfate: 10 µg ml^-1^ and amphotericin B: 25 µg ml^-1^ (Thermo Fisher Scientific), following Zimmer and Bartholme (71). The terraria, sand, and litter were replaced weekly to maintain hygiene.

The animal fitness was followed for 42 days by aseptically measuring their weights (±0.01 g). During feeding, three fresh faeces pellets (0.15–0.19 g for *E. pulchripes* and 0.01–0.02 g for *G. connexa*) were sampled from the terraria, suspended in phosphate buffer (2 ml; pH 7.4), plated in triplicates on LB-agar plates and incubated at 25 **°**C. After 16 h, the colonies were counted and used to quantify the bacterial load. The remaining faecal material was kept at -20 **°**C for DNA extraction (see below). Methane emission was also monitored (see below).

### Inhibition of methanogenesis

Thirty *E. pulchripes* individuals were divided into three groups of ten. The Control group was fed on untreated litter, while the other two groups were fed litter treated with 5 mM or 10 mM of Sodium 2-bromoethanesulfonate (Na-BES; Sigma-Aldrich) to inhibit methanogenesis. Moisture was maintained by spraying with sterile tap water or Na-BES solution every other day. The animals’ weight and CH_4_ production were regularly monitored for 64 days. Methane emission measurements were conducted by placing the millipedes in sealed glass bottles with wet filter paper pieces to maintain humidity (130 ml bottle for *E. pulchripes*; 30 ml for *G. connexa*; Thermo Fisher Scientific) for 4 h at 20 **°**C. The control was glass vessels without animals. Headspace samples (0.5 ml) were collected at the start and the end of incubation using a gas-tight syringe and analysed on a gas chromatograph (HP 5890 series II; Hewlett Packard) equipped with a 2 m Porapak N column at 75 °C and an FID detector. The difference in CH_4_ concentration between start and finish was used to calculate the production rate.

### Identification and enumeration of protists and symbiotic methanogens

Fourteen days post-CH_4_-inhibition, fresh *E. pulchripes* faecal pellets were crushed using a sterilised mortar and pestle, vortexed in 5 ml of 1X phosphate buffer saline (PBS) solution (pH 7.2), and then incubated at room temperature for 2–6 h to dissolve the aggregates. After spin-down, 2 µl of the supernatant was examined under a bright-field microscope (20x) using a Neubauer chamber (Sigma-Aldrich). Protists were identified and enumerated. Part of the supernatant was also used for enumerating the ciliate-associated archaea and methanogens of the *Methanobacteriales* and *Methanomascilliicoccales* orders using Catalysed Reporter Deposition Fluorescence *in situ* Hybridization (CARD-FISH; see Supplementary material for further details).

### Stable isotope labelling of RNA

For the SIP experiment, three replicates from separate terraria were used for each species. *E. pulchripes* had one individual per replicate, while *G. connexa* had five to adjust for size differences. Millipedes were fed 99.9% ^13^C-labelled Canadian-poplar leaves (IsoLife, Netherlands). Control groups were fed unlabelled leaves. Rearing conditions were maintained as described above. Faecal samples were collected every 2 days for isotopic labelling analysis. Then, 1.9 g of faeces from each millipede species were vacuum dried in a SpeedVac DNA130 (Thermo Fisher Scientific) at 45 ℃ for 3 h, and 25 µg was transferred into triplicate tin capsules. Isotopic labelling was assessed at the Stable Isotope Facility, Biology Centre CAS, using a Thermo Scientific^TM^ 253 Plus^TM^ 10 kV IRMS equipped with a SmartEA Isolink and GasBench II (Thermo Fisher Scientific). The ^13^C at% was calculated following Hayes (72; data not shown). Animals were sacrificed and dissected on days 3, 7, 14, and 21 following Sardar *et al.* (23) and stored at -20 °C for subsequent analysis. RNA was extracted from frozen hindgut samples, purified and quantified according to Angel *et al.* (73). Hindgut samples from the SIP experiment measured 0.677–1.108 g for *E. pulchripes* and 0.083–0.092 g for *G. connexa*. See Supplementary material for further details.

### Isopycnic ultra-centrifugation of ^13^C labelled RNA

Following RNA purification, density gradient centrifugation was performed in caesium trifluoroacetate (CsTFA) density gradients following a previously published protocol (74). See Supplementary material for further details.

### Gene quantification, amplicon library construction and sequencing

Pooled faecal pellet samples from the antibiotics curing and inhibition of methanogenesis experiments used for DNA extraction were 0.43–0.59 g for *E. pulchripes* and 0.20–0.40 for *G. connexa.* See Supplementary material for further details. DNA extracts from the antibiotics treatment experiment (24 samples per species) were subjected to 16S-rRNA-gene quantification using the QX200 AutoDG Droplet Digital PCR System (ddPCR; Bio-Rad), primers 338F—805R and the 516P FAM/BHQ1 probe (75). DNA extracts from the methanogenesis inhibition experiment were used for quantifying the *mcrA* gene as a marker for methanogens using primers mlas_mod and mcrA-rev, according to Angel *et al.* (76). Before sequencing, the cDNA from the SIP fractions (160 samples for each millipede species) was used for quantifying the 16S rRNA copies of bacteria using the same method as mentioned above and the 18S rRNA copies of fungi using the FungiQuant system (77). For amplicon sequencing, the V4 region of the 16S rRNA gene was amplified and sequenced in a two-step protocol on an Illumina MiniSeq platform (Mid Output Kit; Illumina) according to Naqib *et al*. (78). PCR amplification was performed on 10 ng of DNA or 2 µl of cDNA with primers 515F_mod and 806R (79), synthesised with the Fluidigm linkers CS1 and CS2 on their 5′ end. Sequencing was performed at the DNA Services Facility at the University of Illinois, Chicago, USA.

### Bioinformatic and statistical analyses

Unless mentioned otherwise, all bioinformatic and statistical analyses were done in R V4.1.1 (80). A linear mixed-effects model (81) was fitted to determine the effect of treatments and time on the millipede weight and microbial load. Differences between treatments in terms of total faecal pellet production, methane emission, *mcrA* and 16S rRNA copies were evaluated using an ANOVA model (82) followed by Tukey’s HSD test for pairwise comparisons (83). Survival analysis of the animals was computed using the Kaplan-Meier estimates (84). Sequencing data was analysed as follows: primer and linker regions were removed from the raw amplicon reads using Cutadapt (V3.5; 85). The raw reads were processed, assembled and filtered using the R package DADA2 (V1.28) with the following non-standard filtering parameters: maxEE = c(2, 2) in the filterAndTrim function and pseudo pooling in the dada function (86). Chimaeras were removed with the removeBimeraDenovo option. The quality-filtered pair-end reads were classified to the genus level using SILVA V138 (87), and those not classified as bacteria or archaea were filtered out. Heuristic decontamination was done using the decontam R package (88), and unique sequences were identified and clustered in an amplicon sequence variant (ASV) table. The resulting tables were imported into the R package Phyloseq (89). Read counts were normalised using median sequencing depth before plotting taxa abundance and after excluding ASVs without taxonomic assignments at the phylum level and those below a 5% prevalence threshold. Alpha diversity indices were computed using the vegan package on unfiltered and non-normalised data (90) and evaluated using the Kruskal-Wallis test (91) and Dunn’s test (92). Corrections for multiple testing were made using the Benjamini-Hochberg (BH; 93) method. Values were compared and converted to a compact letter using the cldList function in the rcompanion package (94). Beta diversity was calculated with a constrained analysis of principal coordinates (CAP; 95). Lastly, a permutational multivariate ANOVA (96); function vegan::adonis) was conducted using the Bray-Curtis distance matrix and the pairwise.adonis2 function (97) to assess combined treatment and pairwise effects on the microbial community.

Differentially abundant genera were identified after sterile feeding or antibiotic treatment using ANCOM-BC2 (98) after removing all ASVs not present in at least two samples or with an abundance of less than 2. Only genera with adjusted P-values ≤ 0.05 and those passing the pseudo-count-addition sensitivity analysis were plotted.

Identification of isotopically labelled ASVs in the SIP experiment using differential abundance analysis followed Angel (99). After initial processing as described above, rare taxa (with <100 total reads, present in <2 SIP fractions in a given gradient and its unlabelled counterpart). The DADA2 output sequences were aligned using sina 1.7.2 (100) against the SILVA V138 DB, and a maximum-likelihood phylogenetic tree was constructed using IQ-TREE V2.1.1 (101) with the ‘-fast‘ option. The 16S rRNA copies were plotted against the density and used to calculate absolute ASV abundances. Fractions with densities >1.795 g ml^-1^ (’heavy’ fractions) from each labelled sample at each time point were compared against their unlabelled counterparts using DESeq2 V1.40.1 (102), using the parametric fit type and the Wald significance test. Log_2_ fold change (LFC) shrinkage was applied using the function lfcShrink (103), and the results were filtered to include only ASVs with a positive log_2_ fold change and a p-value <0.1 (one-sided test).

## Acknowledgements

We are grateful for the support of Lucie Faktorová and Eva Petrová in collecting *G. connexa* samples, Lucie Faktorová in maintaining the *E. pulchripes* colony and assisting with millipede dissection, and Eva Petrová for her guidance and assistance in DNA and RNA extractions and quantification. We are thankful to Radka Malá for her assistance in the filtration and fixation of CARD-FISH samples. Special thanks to Travis Blake Meador, Stanislav Jabinski, and Poláková Ljubov for their contributions to stable isotope detection and quantification in millipede faeces. RA, SG and JEN were supported by a Junior Grant from the Czech Science Foundation (GA ČR), grant number 19-24309Y. The funders had no role in study design, data collection and interpretation, or the decision to submit the work for publication

## Author Contributions

The approach for this study was conceptualised by RA and VS, experiments were carried out by SG, JEN, MMS and TH, and the data analysis was designed by RA and JEN. The bioinformatics analyses were carried out by JEN and RA. The manuscript was written by JEN, SG and RA, with significant contributions from MMS and VS. All authors have thoroughly reviewed and approved the final version of the manuscript.

### Availability of data and analysis scripts

The short-read amplicon sequencing data have been deposited under the NCBI BioProject PRJNA948469 with BioSample SUB13838396 for antibiotics treatment and SUB13843680 for RNA-SIP. For reproducibility, reusability, and transparency, the scripts used in this study are available on GitHub (https://github.com/ISBB-anaerobic/Active-microbial-community-pre-and-post-inhibition.git).

