## Supplementary methods and figure for "Cellulose fermentation by the gut microbiota is likely not essential for the nutrition of millipedes"

Running title: **Gut microbiota in millipede nutrition**

Julius Eyiuche Nweze[a,b], Shruti Gupta[a*], Michaela M. Salcher[a], Vladimír Šustr[a], Terézia Horváthová[c*], Roey Angel[a,b #]

[a]Institute of Soil Biology and Biogeochemistry, Biology Centre CAS, České Budějovice, Czechia

[b]Faculty of Science, University of South Bohemia in České Budějovice, Czechia

[c]Institute of Hydrobiology, Biology Centre CAS, České Budějovice, Czechia

### Supplementary material

### Supplementary methods

#### Identification and enumeration of protists and symbiotic methanogens

In parallel to enumerating the protists using bright-field microscopy, the faecal pellet suspensions were also fixed at 4 °C for 1.5 hours with 2% paraformaldehyde (PFA; Sigma-Aldrich) and subjected to sequential vacuum filtration through 10 µm and 0.2 µm white polycarbonate filters (Sigma-Aldrich). The filters were air-dried and stored at -20 °C for Catalysed Reporter Deposition Fluorescence *in situ* Hybridization (CARD-FISH) analysis. For CARD-FISH, specific horseradish peroxidase (HRP) rRNA-targeting oligonucleotide probes were used (biomers.net). These included a universal probe for archaea (ARC915; Stahl 1991) and specific probes targeting the order *Methanobacteriales* (MB311; Crocetti *et al.* 2006) and *Methanomascilliicoccales* (RC281r_mod; modified from (Iino *et al.* 2013). A nonsense probe (NON-EUB338; (Wallner *et al.* 1993)) served as a negative control (Table S1). Probe coverage and specificity were assessed with TestProbe on ARB Silva (Quast *et al.* 2013). Hybridisation stringency was evaluated *in silico* using the Mathfish platform (Yilmaz *et al.,* 2011) and confirmed with optimised formamide concentration (Sigma-Aldrich; Table S1). The filters were prepared following Pernthaler *et al.* (2002). See Supplementary material for further details. Samples were visualised using an OLYMPUS BX53 (Olympus Optical Ltd.) or a Zeiss Imager Z2 (Carl Zeiss) epifluorescence microscope at 1000X magnification. Methanogens per ciliate were manually counted. Positive (using a general archaeal probe) and negative (no probe and a nonsense probe) control filters were also analysed.

The polycarbonate filters were embedded in warm, 0.2% low-melting agarose (Thermo Fisher Scientific) and dried at 37 °C in an oven. Cells on the filters were permeabilised with a lysozyme solution (10 mg $ml^{-1}$; Sigma-Aldrich) at 37 °C for 45 minutes, then incubated in achromopeptidase solution (2 µl of 30 KU achromopeptidase in 1 ml of NaCl Tris buffer; Sigma-Aldrich) for 15 minutes at 37 °C. For hybridisation, filters were cut, labelled, and hybridised using 300 µl hybridisation buffer and 2 µl probe for 2 hours at 35 °C, followed by 20-30 min wash in a 37 °C washing buffer and incubation in phosphate-buffered saline with Tween-20 for 45 minutes at 37 °C. The signal from the probe was amplified by incubating blot-paper-dabbed filters with fluorescently labelled tyramide (Sigma-Aldrich) and 0.15% $H_2O_2$ for 30 min in the dark. Fluorescein isothiocyanate (FITC; Sigma-Aldrich) was used as a fluorochrome, and 4′,6-diamidino-2-phenylindole (DAPI; Sigma-Aldrich) as a sample counterstain in a mounting solution containing the antifading reagent vectashield.

Samples were counted on an OLYMPUS BX53 (Olympus Optical Ltd.) epifluorescence microscope at a magnification of 1000X with filter sets for DAPI, FITC and autofluorescence. Images were taken with a Zeiss Imager.Z2 (Carl Zeiss) epifluorescence microscope system equipped with an Axiocam 506 (Carl Zeiss), and a Colibri LED light system and filter sets for DAPI (LED module 385 nm; filter set 49), fluorescein (LED module 475 nm; filter set 38 HE) and autofluorescence (LED module 567 nm; filter set 62 HE). Z-stacks (7-11 multi-channel z-stacks) were merged into one image by orthogonal projection with the software ZEN 2.6 (Carl Zeiss).

#### Nucleic acid extraction and quantification

Nucleic acids were extracted as follows: the samples were subjected to three consecutive bead beating rounds (Lysing Matrix E tubes; MP Biomedicals™) in a FastPrep-24™ 5G (MP Biomedicals™) in the presence of CTAB, phosphate buffer (pH 8.0) and TE-buffer-saturated phenol, followed by phenol-chloroform-isoamyl alcohol (25:24:1; all from Carl Roth) purification, precipitation using Invitrogen™ UltraPure™ Glycogen (Thermo Fisher Scientific) and then purification with the OneStep™ PCR Inhibitor Removal Kit (Zymo Research). The resulting DNA was then quantified using the Quant-it™ PicoGreen DNA Assay Kit (Thermo Fisher Scientific). The RNA was purified from the DNA/RNA hindgut extract using TURBO™ DNase and GeneJET RNA Cleanup and Concentration Micro Kit (Thermo Fisher Scientific) for the SIP experiment. The quantity and quality of the RNA were determined using Quant-it™ RiboGreen RNA Assay Kit (Thermo Fisher Scientific) and the Agilent 2100 Bioanalyzer (Agilent Technologies).

#### Isopycnic ultra-centrifugation of $^{13}C$ labelled RNA

Each gradient was prepared with 4.848 ml of CsFTA solution (GE Healthcare), 1.083 ml of gradient buffer (0.1 M Tris-HCl, 0.1 M KCl, 1 mM EDTA; Sigma Aldrich) and 211 µl of Hi-Di Formamide (3.56 % v/v; Thermo Fisher Scientific). The density was confirmed using an AR200 Automatic Digital Refractometer (Reichert) against a calibration curve, and occasionally, adjustments were made using CsTFA or gradient buffer until a final density of 1.79 g $ml^{-1}$ was reached. Finally, each 6 ml Ultracrimp PA centrifugation tube (Thermo Fisher Scientific) contained approximately 5.8 ml of the density gradient solution and ca. 500 ng of RNA. Additionally, a control tube containing no RNA was used to exclude the presence of DNA or RNA contamination. Tubes were centrifuged in a TV-1665 vertical rotor in a Sorvall WX Ultra 100 Ultracentrifuge (Thermo Fisher Scientific) at 20 °C and 130,000 ×g for 72 h. Twelve fractions of ca. 500 µl were collected into 2.0 ml low-binding collection tubes (Eppendorf) using a NE-300 Just Infusion™ Syringe Pump (NEW ERA PumpSystem Inc.), and the buoyant density (BD) of each fraction from the control tube was determined using a refractometer. RNA was then precipitated from the gradient fractions using 2 µl of GlycoBlue (Thermo Fisher Scientific), 47 µl of 3M Na-Acetate (pH 5.5) (Thermo Fisher Scientific) and 1.175 ml ethanol (absolute), and dissolved in 10 µl of RNA Storage solution (Thermo Fisher Scientific). Lastly, RNA from the gradient fractions 2–11 was converted into cDNA in 20 µl reactions using SuperScript IV RT (Thermo Fisher Scientific). The resulting cDNA was then stored at −20 °C until further processing.

### Supplementary figures

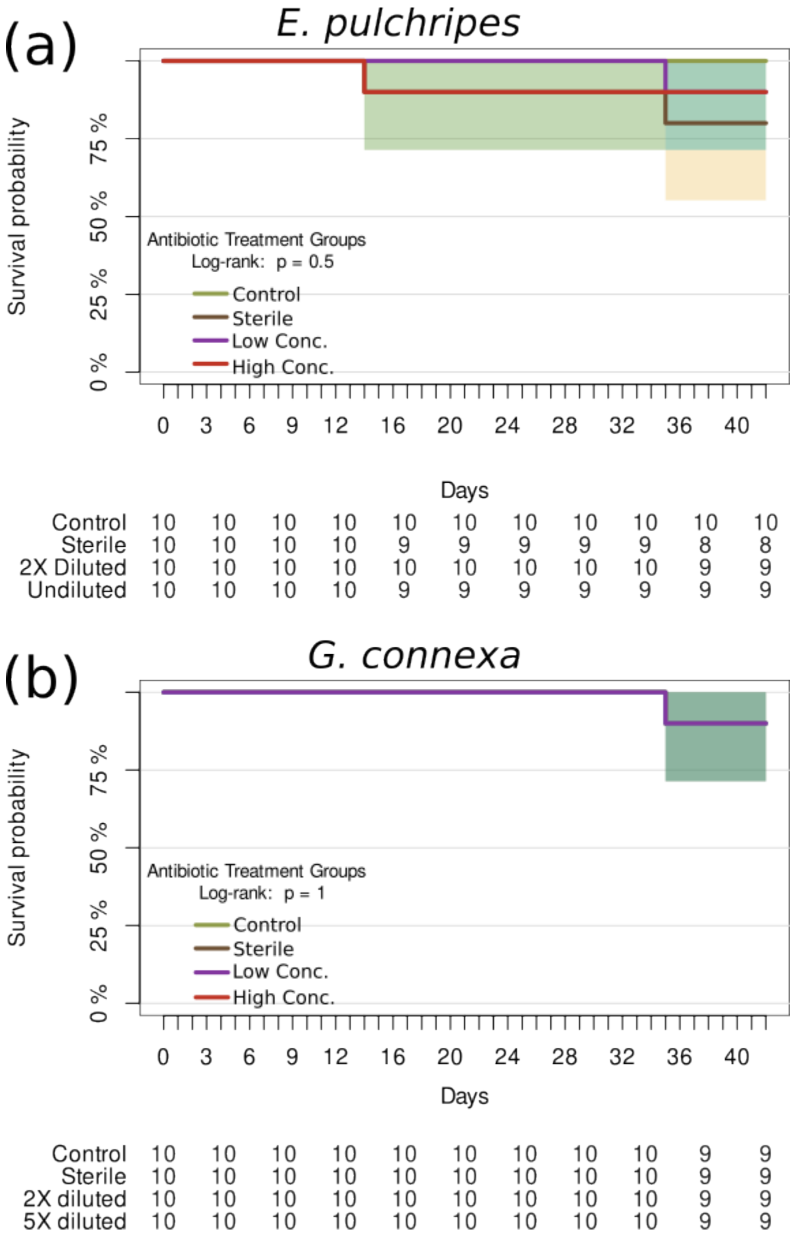


**Fig. S1 Kaplan-Meier survival curves based on treatment for *E., pulchripes* and *G., connexa*.** The study involved a total of 40 individuals, evenly distributed across four groups. 'High Conc.' and 'Low Conc.' refer to the concentration of applied antibiotics (see Materials and Methods for more details).

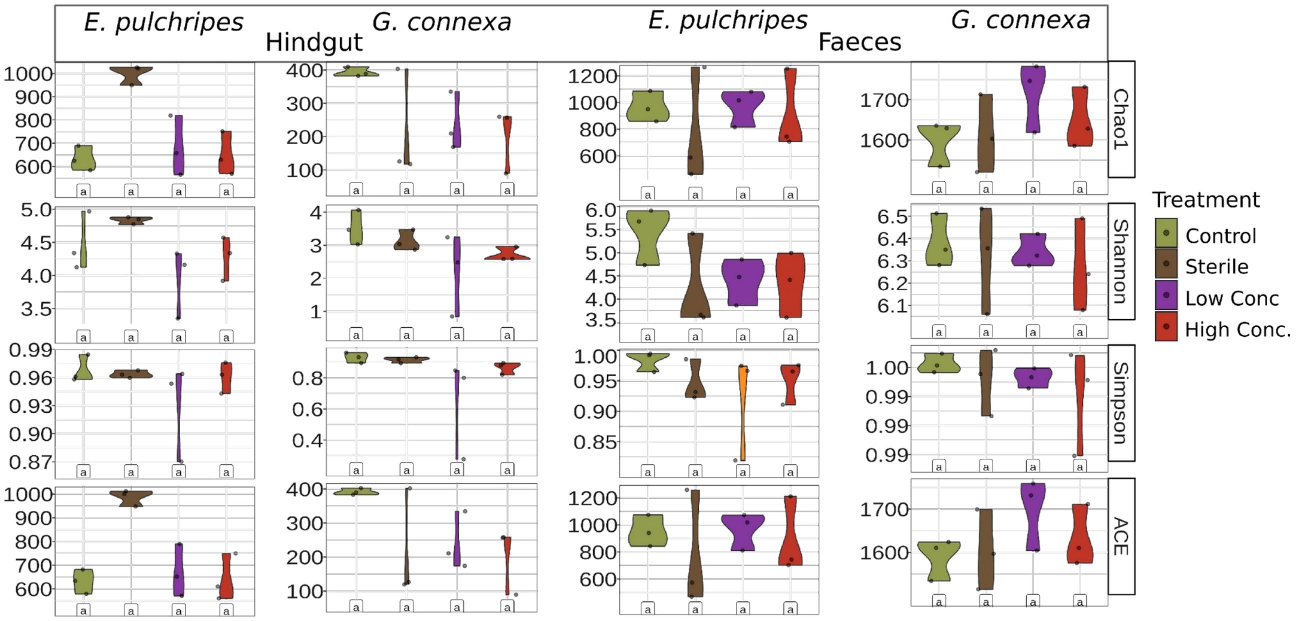


**Fig. S2 Alpha diversity indices of the microbial communities in the hindgut and faeces after antibiotics treatment in *E. pulchripes* and *G. connexa*.** Additional alpha diversity values for each species to those shown in Fig. 3, stratified by treatment groups of hindguts and faeces from *E. pulchripes* and *G. connexa*. The statistical test was based on Kruskal–Wallis (identical letters indicate p > 0.05). ‘High Conc.' and 'Low Conc.' refer to the concentration of applied antibiotics (see Materials and Methods for more details).

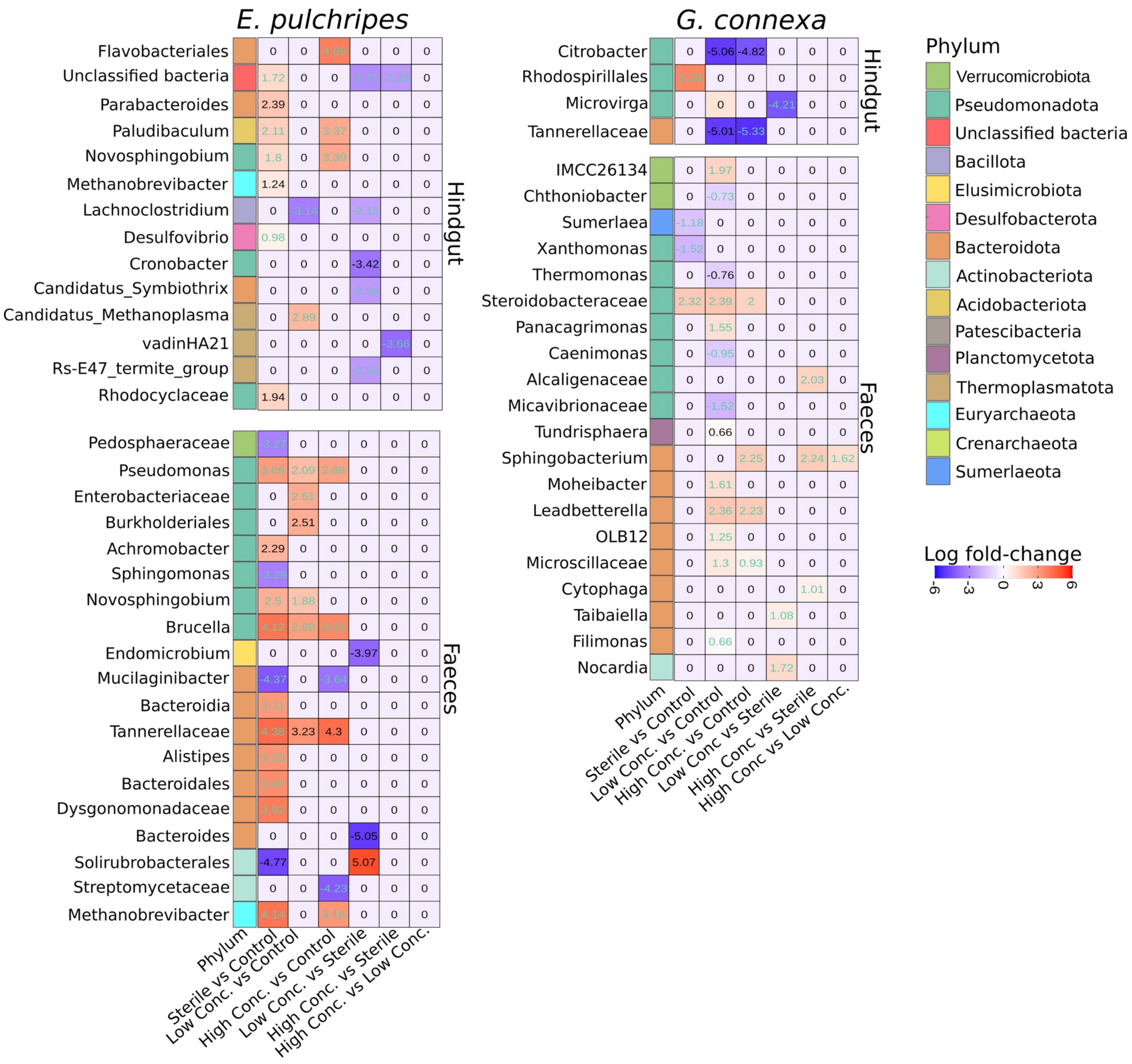


**Fig. S3 Heatmaps of the ANCOM-BC2 pairwise analysis for the effect of antibiotics or sterile feeding on the microbial relative abundances in the hindgut and faecal samples from *E. pulchripes* and *G. connexa*.** The heatmaps show multiple pairwise comparisons on the genus level among the four groups: control, sterile-fed, low-conc. and high-conc. antibiotics. The X-axis represents treatment comparison, while the Y-axis displays significant genera and their phylum, identified by ANCOM-BC2. Each cell is colour-coded, with blue representing reduced abundance and red representing increased abundance in response to the treatment. The numbers in each cell indicate the log fold change. The Benjamini-Hochberg method was used to correct for multiple testing, and taxa with log fold-change values marked in green have successfully passed the sensitivity analysis for pseudo-count addition. Refer to Fig. S1 for details regarding the treated groups.

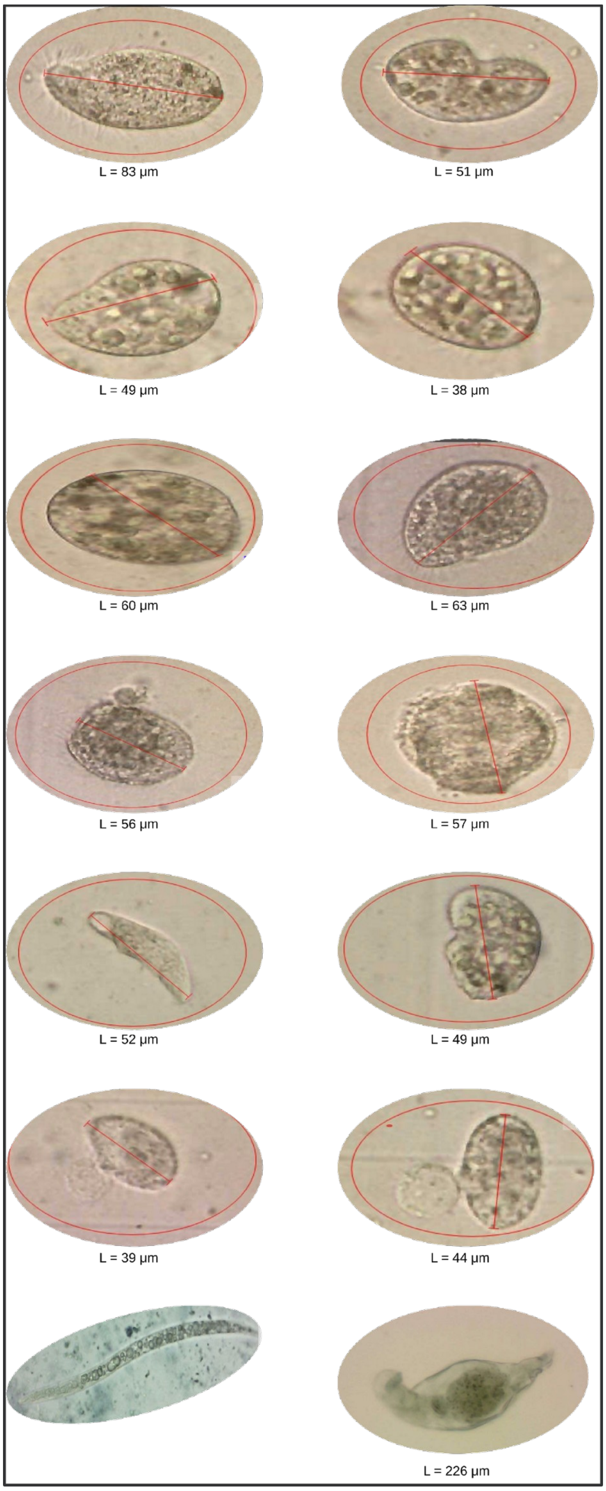


**Fig. S4 Light-microscopy crop images of symbiotic ciliates, nematode and rotifer found in the faeces of *E. pulchripes*.** "L" represents the length of the organisms as measured from the image.

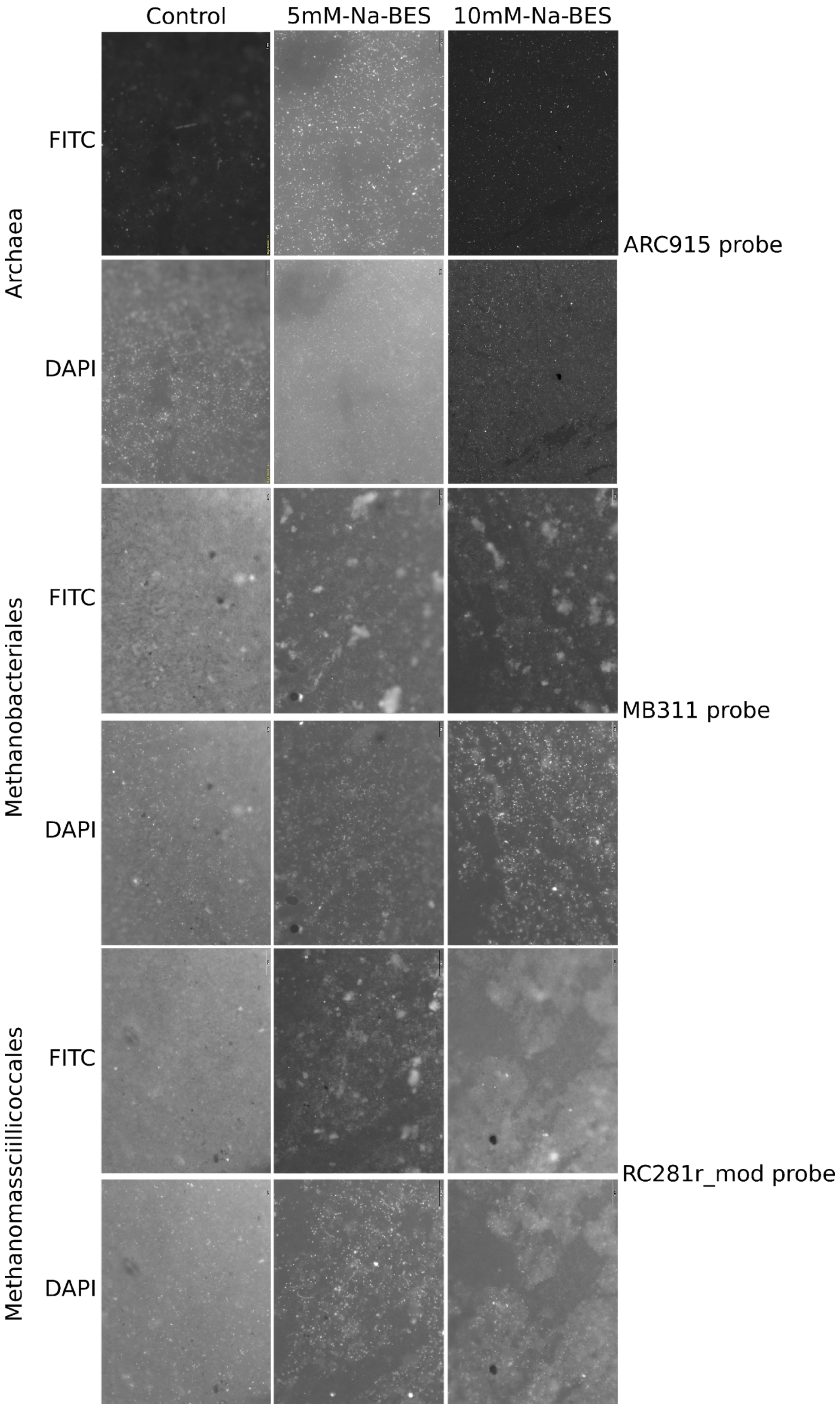


**Fig. S5 CARD-FISH images of filtered cells from faecal samples of Na-BES-fed *E. pulchripes*.** The free-living or detached methanogens were captured from faecal samples using a 0.2 µm filter and labelled with the following probes: ARC915 for general archaea, MB311 for *Methanobacteriales*, and RC281r_mod for *Methanomassciillicoccales*. Each FITC-probe image has a parallel DAPI-stained image. The treatments were represented by control (group fed with untreated litters); 10mM-Na-BES-treated litters (group fed with 10mM-Na-BES-treated litters); and 5mM-Na-BES-treated litters (group fed with 5mM-Na-BES-treated litters).

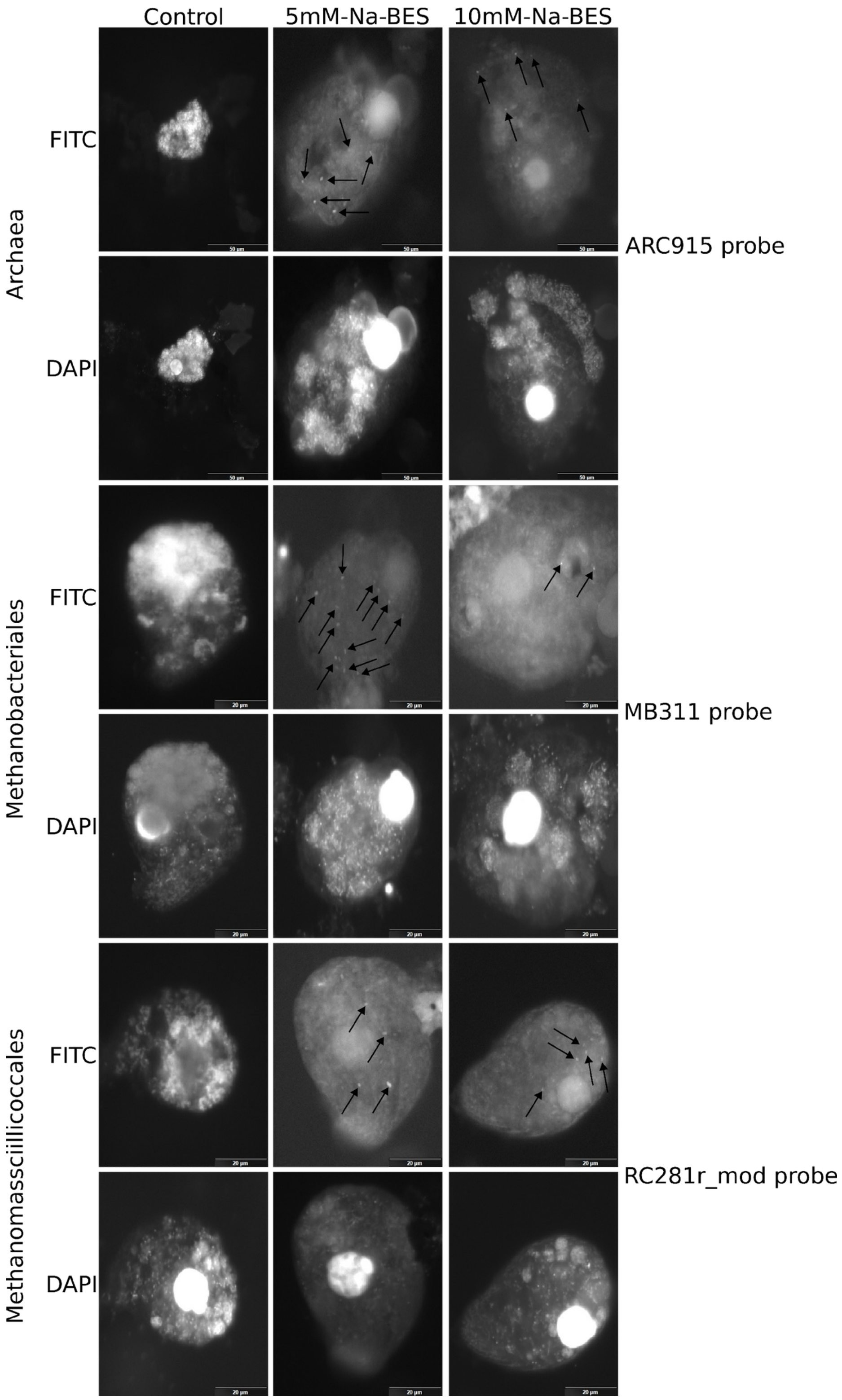


**Fig. S6. CARD-FISH images of ciliate-associated cells from faecal samples of Na-BES-fed *E. pulchripes*.** Ciliates were captured from faecal samples using a 10 µm filter and labelled with the following probes: ARC915 for general archaea, MB311 for *Methanobacteriales*, and RC281r_mod for *Methanomassciillicoccales*. Each FITC-probe image has a parallel DAPI-stained image.

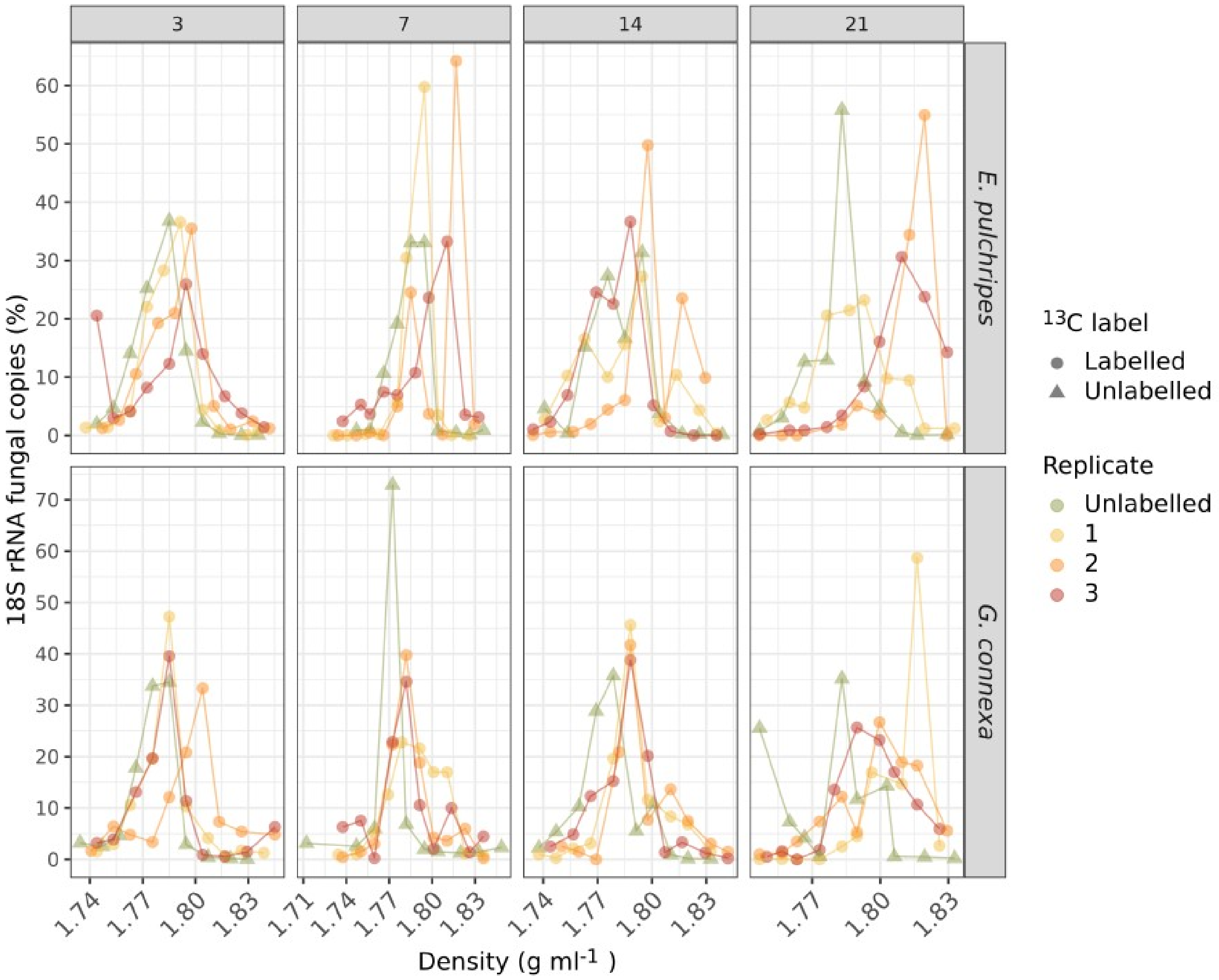


**Fig. S7. Fungal 18S rRNA copies recovered from each fraction in the SIP gradients.** Values on the y-axis are the rRNA copies relative to the total number of rRNA copies obtained from the entire gradient in %. Values on the x-axis show the buoyant density of each fraction. Labelled RNA is expected to be found in fractions with a density >1.795 g ml$^{-1}$.

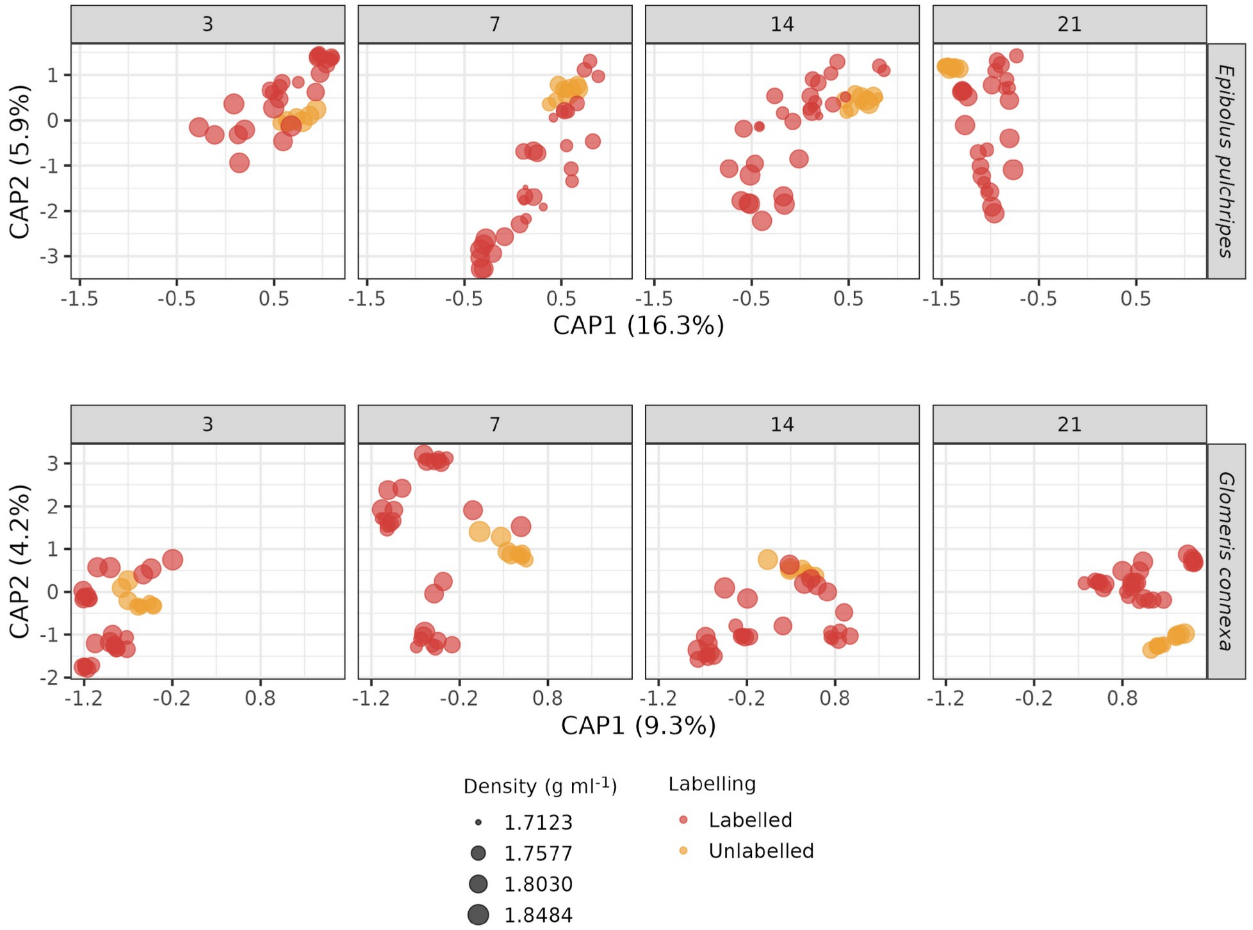


**Fig. S8.** Constrained principal coordinates analysis (PCoA) of Morisita-Horn dissimilarities in community composition of rRNA sequences from the SIP fractions. An ordination model using the formula: Dist.Mat ~ Day + Density.zone was calculated for each millipede species separately.

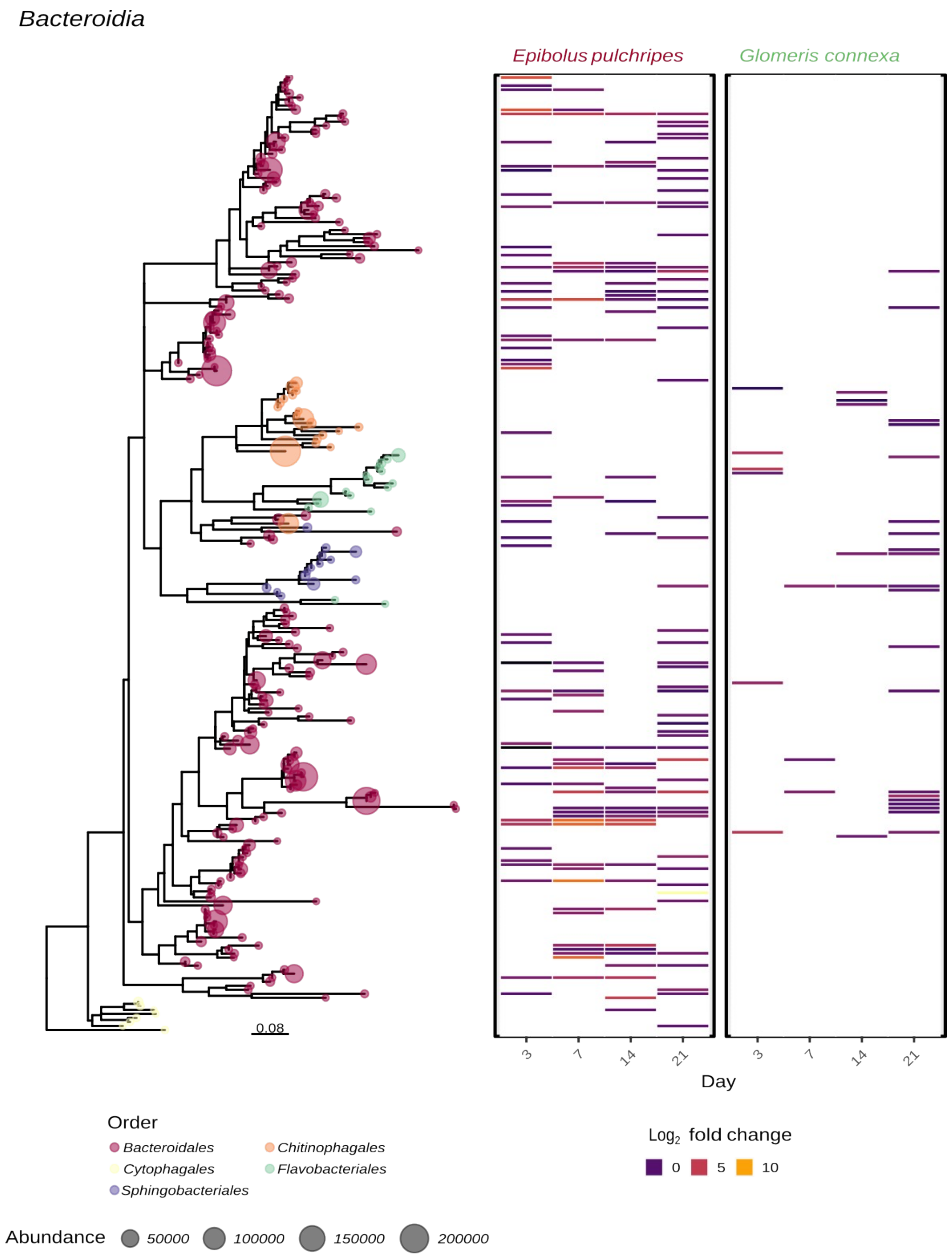


**Fig. S9. A heatmap and a phylogenetic tree of ASVs from the class *Bacteriodia* (phylum: *Bacteroidota*).** Each tip in the tree represents an ASV, and its circle size is proportional to the combined abundance. The tips are also colour-coded according to the order to which they are classified. Each heatmap column represents a time point, and cells of labelled ASVs are filled to represent their $Log_2$-fold change in abundance compared to the unlabelled controls.

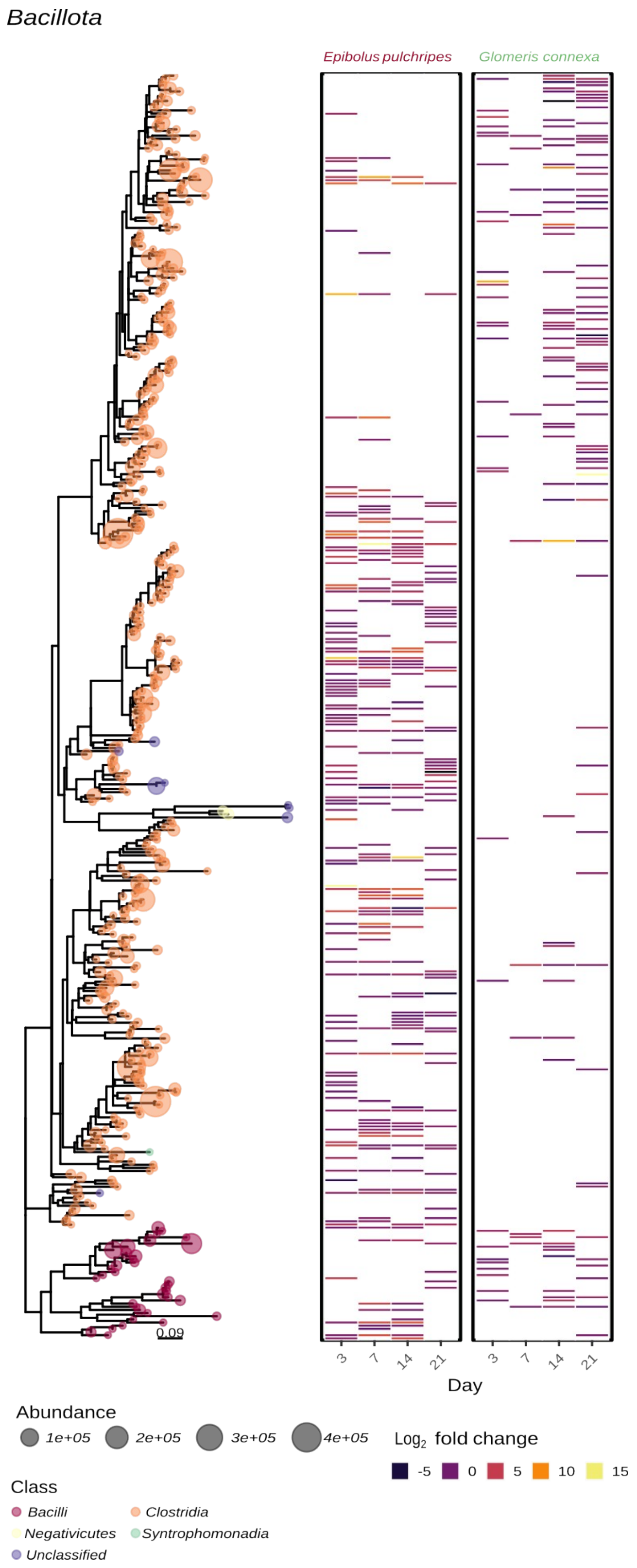


**Fig. S10. A heatmap and a phylogenetic tree of ASVs from the phylum *Bacillota*.** Each tip in the tree represents an ASV, and its circle size is proportional to the combined abundance. The tips are also colour-coded according to the class to which they are classified. Each heatmap column represents a time point, and cells of labelled ASVs are filled to represent their $Log_2$-fold change in abundance compared to the unlabelled controls.

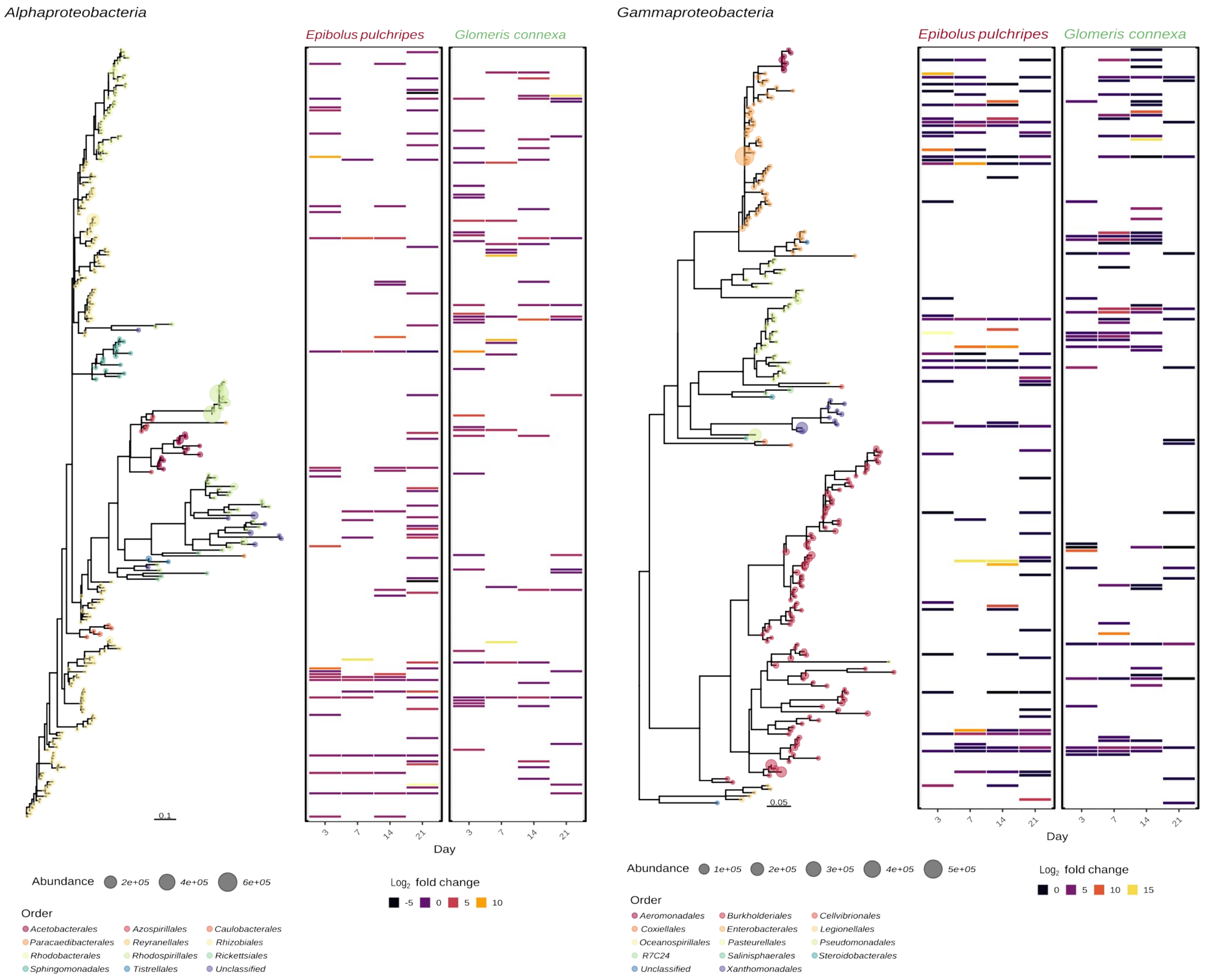


**Fig. S11. A heatmap and a phylogenetic tree of ASVs from the classes *Alphaproteobacteria* and *Gammaproteobacteria* (phylum: *Pseudomonadota*).** Each tip in the tree represents an ASV, and its circle size is proportional to the combined abundance. The tips are also colour-coded according to the order to which they are classified. Each heatmap column represents a time point, and cells of labelled ASVs are filled to represent their $Log_2$-fold change in abundance compared to the unlabelled controls.

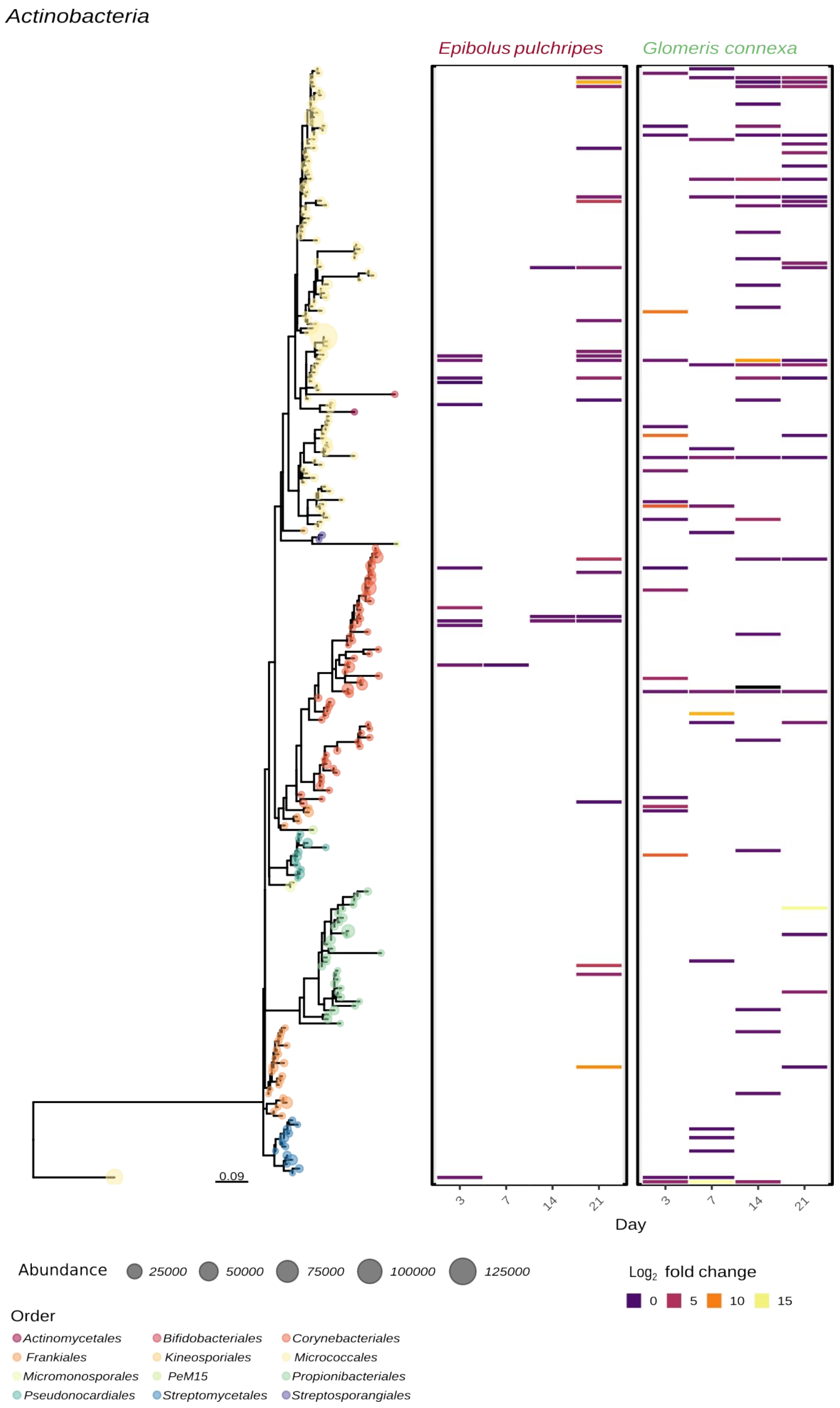


**Fig. S12. A heatmap and a phylogenetic tree of ASVs from the class *Actinobacteria* (*Actinomycetia;* phylum *Actinomycetota*).** Each tip in the tree represents an ASV, and its circle size is proportional to the combined abundance. The tips are also colour-coded according to the order to which they are classified. Each heatmap column represents a time point, and cells of labelled ASVs are filled to represent their $Log_2$-fold change in abundance compared to the unlabelled controls.

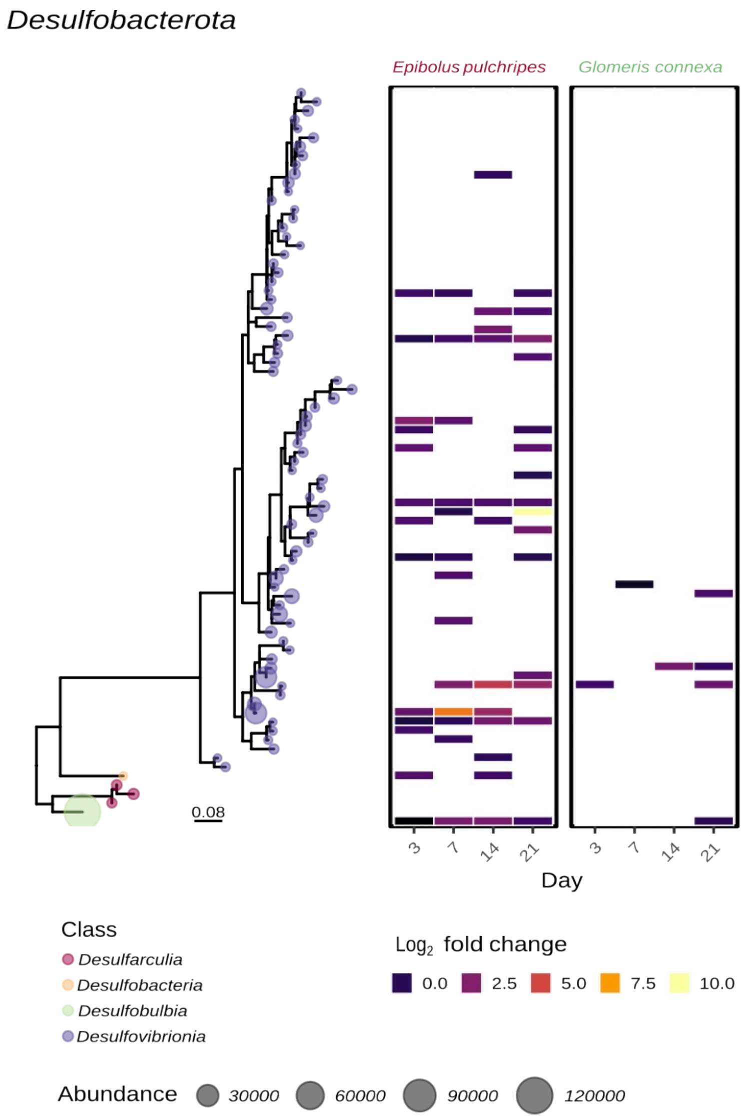


**Fig. S13. A heatmap and a phylogenetic tree of ASVs from the phylum *Desulfobacterota*.** Each tip in the tree represents an ASV, and its circle size is proportional to the combined abundance. The tips are also colour-coded according to the class to which they are classified. Each heatmap column represents a time point, and cells of labelled ASVs are filled to represent their $Log_2$-fold change in abundance compared to the unlabelled controls.

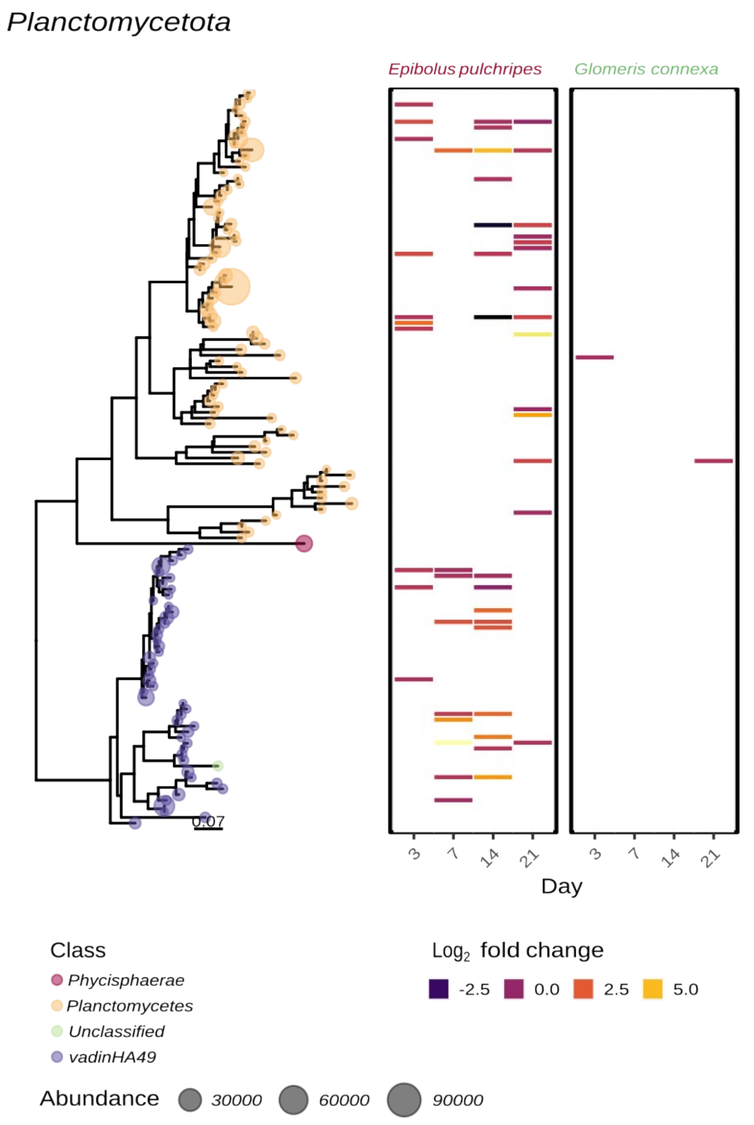


**Fig. S14. A heatmap and a phylogenetic tree of ASVs from the phylum *Planctomycetota*.** Each tip in the tree represents an ASV, and its circle size is proportional to the combined abundance. The tips are also colour-coded according to the class to which they are classified. Each heatmap column represents a time point, and cells of labelled ASVs are filled to represent their $Log_2$-fold change in abundance compared to the unlabelled controls.

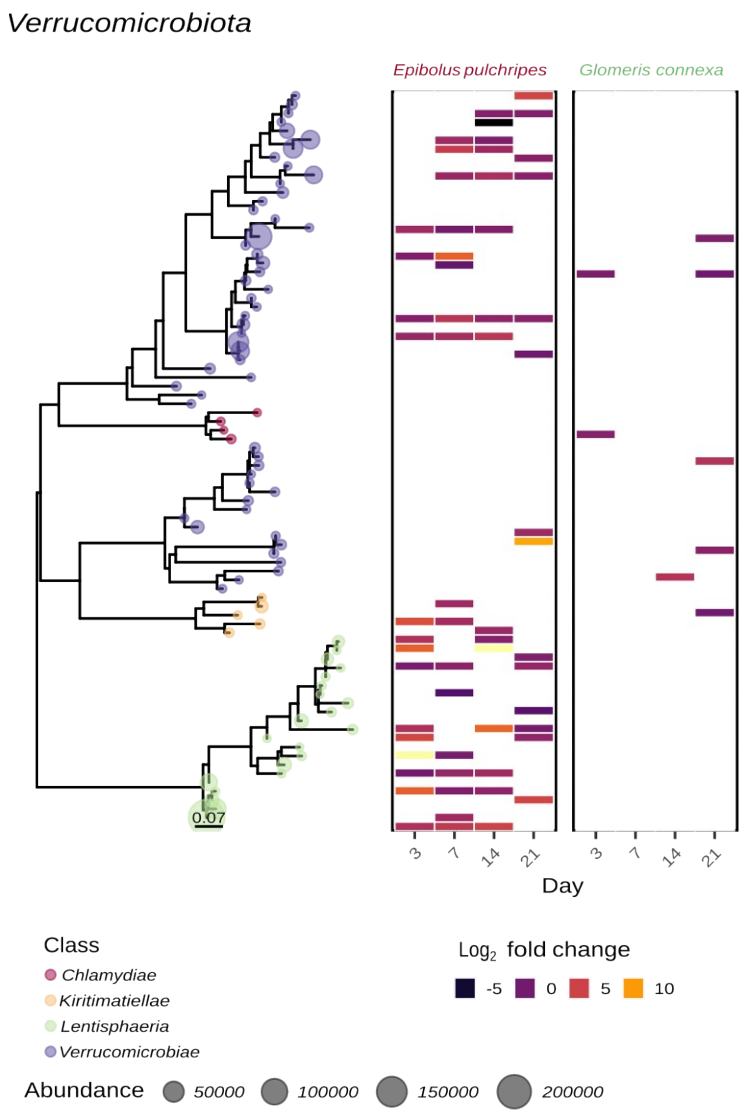


**Fig. S15. A heatmap and a phylogenetic tree of ASVs from the phylum *Verrucomicrobiota*.** Each tip in the tree represents an ASV, and its circle size is proportional to the combined abundance. The tips are also colour-coded according to the class to which they are classified. Each heatmap column represents a time point, and cells of labelled ASVs are filled to represent their $Log_2$-fold change in abundance compared to the unlabelled controls.
